# CORE GREML: Estimating covariance between random effects in linear mixed models for genomic analyses of complex traits

**DOI:** 10.1101/853515

**Authors:** Xuan Zhou, Hae Kyung Im, S. Hong Lee

**Affiliations:** Australian Centre for Precision Health, University of South Australia, Adelaide, South Australia, Australia; Section of Genetic Medicine, Department of Medicine, The University of Chicago, Chicago, Illinois, USA; South Australian Health and Medical Research Institute, Adelaide, South Australia, Australia

**Author notes:** Correspondence: S. Hong Lee.

## Abstract

Linear mixed models (LMMs) using genome-based restricted maximum likelihood (GREML) are a key variance partitioning tool, where effects of multiple sources, such as different functional genomic regions, on phenotypes are treated as random. Classic LMMs assume independence between random effects, which can cause biased estimation of variance components. Here, we relax this independence assumption by introducing a generalised GREML, called CORE GREML, that can explicitly estimate the covariance between random effects. Using extensive simulations, we show that CORE GREML outperforms the conventional GREML, providing unbiased estimates of variance and covariance components. Using data from the UK biobank, we demonstrate that CORE GREML is useful for genomic partitioning analyses and for genome-transcriptome partitioning of phenotypic variance. For example, we found that the transcriptome, imputed using genotype data, explained a significant proportion of phenotypic variance for height (0.15, se = 5.4e-3, *p*-value = 1.5e-283), and that these transcriptomic effects on phenotypes correlated with effects of the genome (r = 0.35, se = 4.6e-2, *p*-value = 1.2e-14). We conclude that the covariance between random effects is a key parameter that needs to be estimated, especially when partitioning phenotypic variance by omic layer.

## Introduction

Genome-wide association studies (GWAS) have been incredibly successful in identifying genetic variants associated with complex traits. However, the proportion of phenotypic variance explained by genome-wide significant single nucleotide polymorphisms (SNPs) is far lower than the narrow-sense heritability estimate^1^. This is largely because GWASs typically examine SNP-trait associations one at a time, generating a large number of tests across the genome for which the stringent Bonferroni correction is applied. To overcome this problem, a whole-genome approach that jointly considers all available SNPs has been introduced^2^, allowing estimation of the proportion of phenotypic variance explained by genome-wide SNPs, i.e. SNP-based heritability. Central to this approach is the use of linear mixed models^3^ (LMMs)—extensions of random-effects models or variance-component models^4^—that treat SNP effects as random.

Using genome-based restricted maximum likelihood (GREML) for parameter estimation, LMMs are a key tool not only for SNP-based heritability of complex traits but also for variance partitioning in general. For example, when heritability is partitioned by functional annotation of SNPs using a LMM with multiple random effects, GREML estimation has provided important insights into the latent genetic architecture of complex traits^5^. As multi-omics data become increasingly available^6^, variance partitioning using LMMs will become indispensable to uncover the relative contributions of multiple ‘omes’ to phenotypic variation. Alongside GREML, linkage disequilibrium score regression (LDSC) provides an alternative way for SNP-based heritability estimation and variance partitioning, using only GWAS summary statistics without the need to access individual genotypes^7,8^. LDSC or stratified LDSC also treats SNP effects as random^8^; and for the same set of individual-level genotype data, this approach generates similar estimates as GREML^9–11^.

Following classic LMMs^3,4^, GREML assumes independence between random effects when estimating variance components. However, it is questionable if this assumption is always valid especially for genomic analyses of complex traits. For example, gene regulatory networks shared between functional categories may generate non-negligible correlations between effects of these categories on phenotypes^12^. In the context of phenotypic variance partitioning by multi-omics layers, effects of genetic variants and their expression levels on phenotypes are likely correlated^13–15^, as exemplified by overlaps between GWAS loci and expression quantitative trait loci (eQTL; e.g.^16,17^). Given these justifiable covariance terms in genomic analyses of complex traits, the naïve assumption of independence between random effects held by GREML can lead to biased partition of phenotypic variance and false inferences on the underlying architecture of complex traits.

Here, we introduce an alternative GREML, referred to as CORE GREML (CORE for COvariance between Random Effects), that fits the Cholesky decomposition of kernel matrices in a LMM to estimate the covariance between a given pair of random effects. Using extensive simulations, we show that CORE GREML outperforms GREML, providing unbiased estimates of variance and covariance components. We also apply CORE GREML to real data to demonstrate its use for genomic partitioning analyses and for genome-transcriptome partitioning of phenotypic variance.

## Results

### Methods overview

The proposed method, CORE GREML, is an extension of GREML, in that it uses the Cholesky theorem to derive the covariance structure between relationship kernel matrices of random effects for mixed-model based partitioning of phenotypic variance (see Methods for details). To validate CORE GREML, we simulated 500 replicates of phenotypic data (n = 10,000) under settings where the covariance between phenotypic effects of random terms was zero, positive and negative (see Supplementary Table 1 for parameter settings). By comparing the model fit by GREML with that by CORE GREML for replicates simulated under the null setting (i.e., no covariance), we estimated the type I error rate of detecting covariance between random effects. Under all settings, we also determined the extent to which CORE GREML recovered the true values of model parameters, and compared CORE GREML and GREML estimates to show the impact of neglecting covariance terms. To facilitate interpretation of results from real data analyses, phenotypic observations under all settings were simulated using available genomic and transcriptomic data as for real data analyses.

For analysis of real data, we selected ten traits with the highest heritability estimates (see Supplementary Figure 1 for SNP heritability estimates) from the UK biobank data that are available to us (reference number 14575), including standing height, sitting height, body mass index (BMI), heel bone mineral density, fluid intelligence, weight, waist circumference, hip circumference, diastolic blood pressure, and years of education. For each trait, we conducted two separate sets of variance partitioning analyses, genomic partitioning by functional region and genome-transcriptome partitioning of phenotypic variance. For each analysis, we applied GREML and CORE GREML and compared the model-fit of the two methods to test the significance of the covariance terms between random effects. Where necessary, we performed a 5-fold cross-validation to compare the prediction accuracy of CORE GREML against that of GREML. Of note, both GREML and CORE GREML use relationship kernel matrices for variance-components estimation (Methods). The kernel matrices for genomic partitioning analyses were constructed using genotypes of 75,396 SNPs from coding regions, untranslated regions and promotors (collectively referred to as ‘regulatory regions’ thereafter), 255,665 from the DNase I hypersensitivity sites (DHS), and 799,935 for all other regions (referred to as ‘other regions’ thereafter). For the genome-transcriptome partitioning of phenotypic variance, the kernel matrix for genetic variance estimation was constructed using genotypes of 1,133,273 genome-wide SNPs, and the kernel matrix for the estimation of phenotypic variance explained by the transcriptome was based on imputed expression levels of 227,664 genes from 43 tissues^18^ (see Supplementary Table 2), respectively. Importantly, our primary interest was not variance partitioning per se; rather our intention was to demonstrate the use of CORE GREML to detect and estimate covariance terms between random effects in mixed-model based variance component analyses and to show the impact of neglecting covariance terms on variance-components estimation.

### Method validation by simulation

For phenotypic data simulated under the genomic partitioning model (see Methods) with zero covariance between effects of genomic regions on phenotypes (i.e., the null setting), the CORE GREML vs. GREML comparison yielded significant results for 19 replicates out of 500, giving an estimated type I error rate of 0.038 for detecting covariance terms. Similarly, for data simulated using the genome-transcriptome model under the null setting (see Methods for the simulation model), the estimated type I error rate was 0.042. Thus, for both simulation scenarios, type I error rate was not inflated.

In terms of parameter estimation, regardless of the simulation model and the parameter setting, CORE GREML consistently yielded unbiased estimates of all model parameters (Supplementary Figures 2 & 3). In contrast, GREML only produced unbiased variance estimates of random effects under the null setting where random effects are not correlated (cov = 0 in Supplementary Figures 2 & 3). As expected, GREML overestimated and underestimated the variance of random effects in the presence of positive and negative correlations between random effects, respectively, and the biased estimation was evident for both genomic partitioning and genome-transcriptome partitioning of phenotypic variance (Supplementary Figures 2 & 3). Thus, our simulation results validate that CORE GREML properly partitions phenotypic variance whether or not the random effects in a LMM are correlated with each other. These results also indicate that GREML would produce biased variance-components estimates when random effects are correlated.

### Assumption on genetic architecture

Incorrect assumptions in the estimation model about the genetic architecture of the trait can also bias variance-components estimation in the context of GREML (e.g.^19^). Therefore, we tested the extent to which CORE GREML estimation is sensitive to a wrong assumption of the genetic architecture in the estimation model. This was achieved by simulating phenotypes under different genetic architectures (see Methods) and comparing CORE GREML estimation from fitting an estimation model that has the correct assumption about the genetic architecture, referred to as the ‘true model’, with that from fitting a ‘wrong model’ that has an incorrect assumption about the genetic architecture.

We found that misspecification of genetic architecture in the estimation model in general biased CORE GREML estimation of variance components but not the covariance term for genome-transcriptome partitioning of phenotypic variance (see Supplementary Figure 4). Nonetheless, under any given genetic architecture, misspecification can be feasibly diagnosed by comparing the likelihood of estimation models that assume a wide range of possible genetic architectures. As shown in Supplementary Figure 5, differences in the likelihood of estimation models are highly indicative of deviations from the true underlying genetic architecture. In fact, a grid search approach has been practiced choosing the model closest to the true underlying genetic architecture in the GREML context^19^.

In light of the above results, to reduce the chance of misspecification of genetic architecture for real data analyses, we fitted two estimation models, the Genome-wide Complex Trait Analysis (GCTA) model^2^ and the Linkage Disequilibrium Adjusted Kinships (LDAK) model (with parameter α, which controls the extent to which minor allele frequency affects the variance of SNP-specific effects on phenotypes, set at the recommended default −0.25^19^). We found that for all traits, the GCTA model had a better fit than the LDAK model, irrespective of estimation method (i.e., GREML or CORE GREML; see Supplementary Tables 3 & 4), indicating that the GCTA model is closer to the true genetic architecture than the LDAK model for our selected traits. Nonetheless, heritability estimates by the two models do not differ substantially (Supplementary Table 3), and significant covariance terms detected by the GCTA model remain significant when using the LDAK model (Supplementary Table 4; although the GCTA model seems more conservative than the LDAK model for detecting covariance terms). This is consistent with the previous observation that heritability estimates based on high-quality common SNPs are robust to variation in assumed genetic architecture (more specifically, parameter α values of the LDAK model; see Figure 6 & Supplementary Figure 4 in Speed et al.^19^). Given the above, unless specified otherwise, results presented in the main text below are GCTA-based; LDAK-based results are included in supplementary materials.

### Real data analyses

Intuitively, the covariance between any pair of random effects would not exist if the variance of any of the random effects is negligible. Therefore, prior to covariance estimation, we tested if the variance components of interest differ from zero for the ten selected traits. For genomic partitioning, we estimated genetic variance by functional region using GREML and found that for all traits all variance components were different from zero by Wald tests (Supplementary Table 5). For genomed-transcriptome partitioning of phenotypic variance, given that all selected traits are high in heritability (i.e., large genetic variances), we tested if the imputed transcriptome could explain a significant proportion of phenotypic variance. To do so, we fitted two models using GREML, a ‘G model’ that breaks phenotypic effects into effects of the genome and residuals, i.e., y=g+ε, and a ‘G-T model’ that decomposes phenotypic effects into effects of the genome and the imputed transcriptome and residuals, i.e., y=g+t+ε. We declared the presence of phenotypic effects of the imputed transcriptome when the G-T model had a better fit than the G model using the likelihood ratio test with one degree of freedom. We found significant phenotypic effects of the imputed transcriptome for all traits except fluid intelligence (Figure 1). Interestingly, although the G-T model had a much better fit than the G model for the nine traits, the two models explained a similar amount of phenotypic variance (Figure 1), which was verified by additional simulations (see Supplementary Note). This suggests that the partition of phenotypic variance represented by the G-T model is closer to the true underlying model than that represented by the G model.

**Figure 1.**
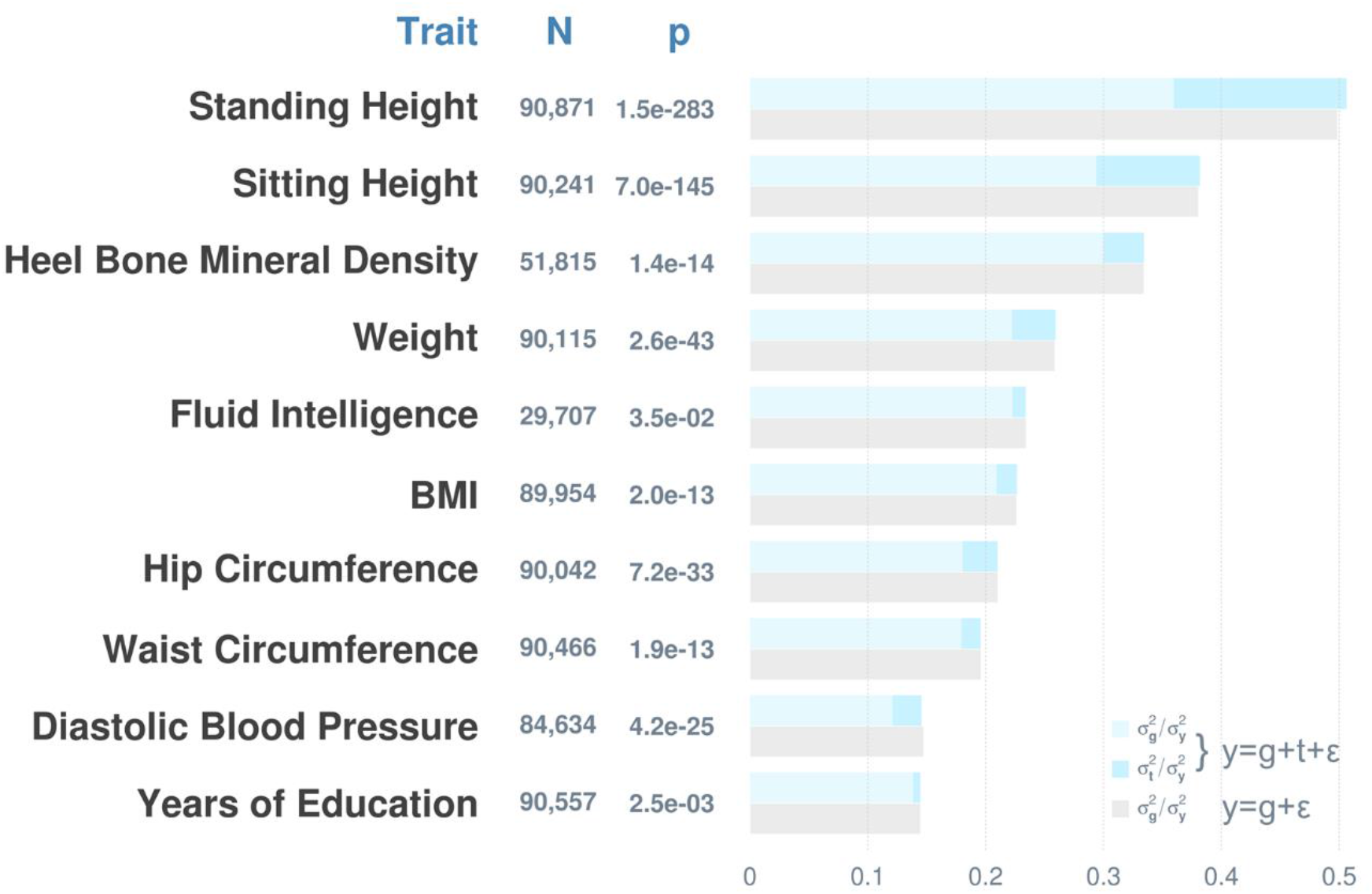
Imputed transcriptome contributes to phenotypic variance. Shown are estimated variance components as a proportion of total phenotypic variance from a linear mixed model that includes a random term for the phenotypic effects of the imputed transcriptome and another model that does not, denoted as y = g+t+ε and y = g+ε, respectively. P-values are from likelihood ratio tests (df = 1) that compare the two models for detecting phenotypic effects of the imputed transcriptome. g = phenotypic effects of the genome; t = phenotypic effects of the imputed transcriptome; ε = residuals; 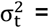 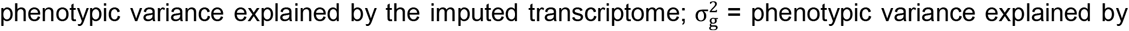 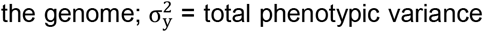. The imputed transcriptome consists of expression levels of 227,664 genes from 43 non-sex specific tissues.

To validate the transcriptomic effects on phenotypes revealed by the G-T model, we performed a 5-fold cross-validation, in which the phenotypic prediction accuracy of the G-T model was compared against that of the G model. For each trait, we randomly split the sample into a training set (~80%) and a validation set (~20%) and iterated this process five times in a manner such that validation sets did not overlap across iterations. To derive the prediction accuracy for the two models, we computed the Pearson’s correlation coefficient between the observed and predicted phenotypes of each trait in each iteration and averaged correlation estimates across five iterations. Figure 2 a. shows that the gain in the phenotypic prediction accuracy by the G-T model relative to that by the G model grew as the estimated transcriptomic contribution to phenotypic variance increased (*p* = 1.86e-06), suggesting that the transcriptomic effects on phenotypes of the selected traits are genuine. Taken together, our results thus far indicate that the variance components of interest differ from zero for all traits with fluid intelligence being the exception for the genome-transcriptome partitioning of phenotypic variance. Importantly, these results established the basis for our subsequent covariance estimation.

**Figure 2.**
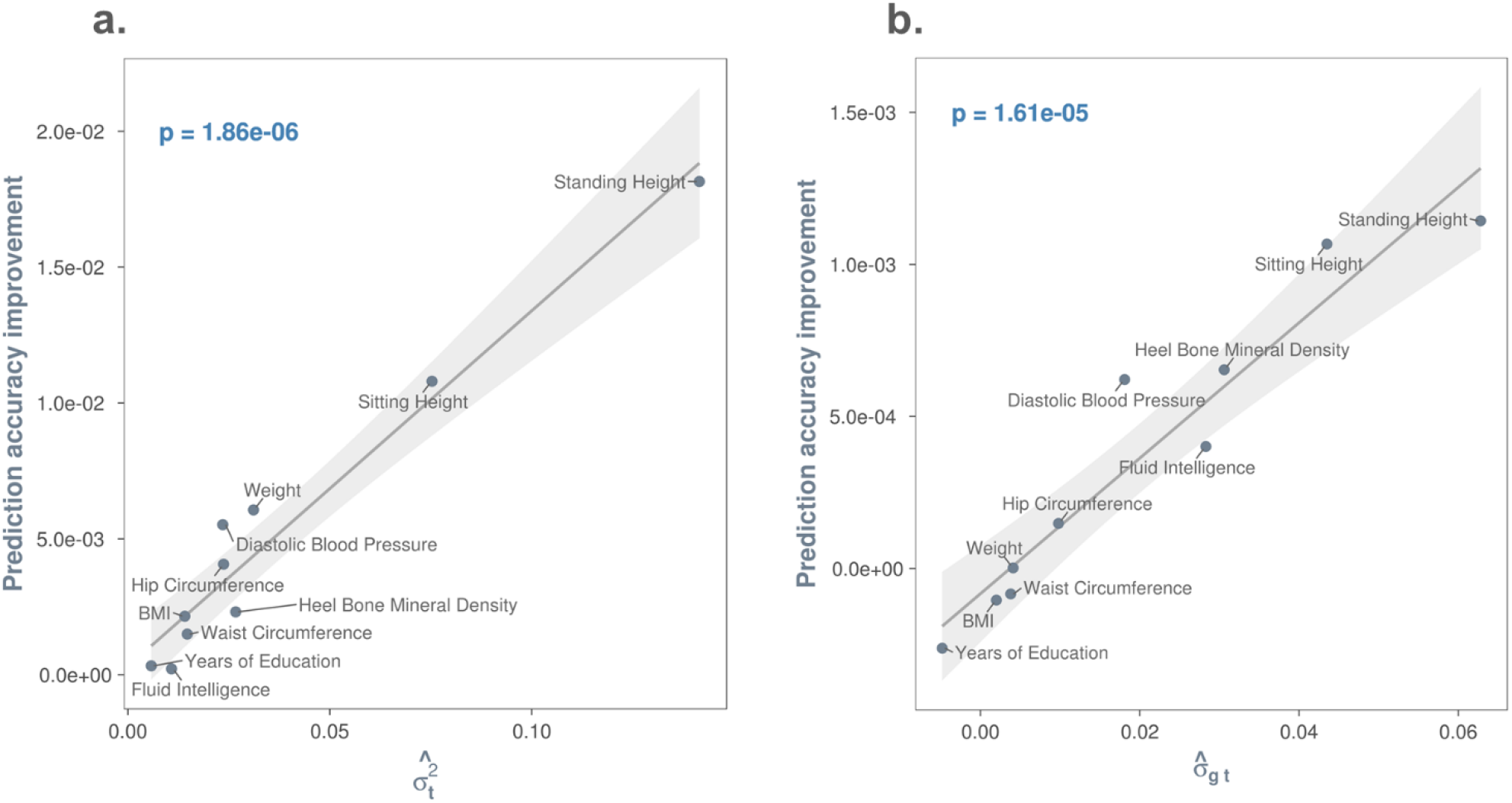
Imputed transcriptome and covariance between phenotypic effects of the genome and of the imputed transcriptome improve phenotypic prediction accuracy. **Panel a.** the more the imputed transcriptome contributes to phenotypic variance, the greater the gain in phenotypic prediction accuracy. 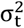 denotes the phenotypic variance explained by the imputed transcriptome. Prediction accuracy was computed using the Pearson’s correlation coefficient between the observed and the predicted for models with and without a random term for phenotypic effects of the imputed transcriptome, denoted as y = g+t+ε and y = g+ε, respectively. g = phenotypic effects of the genome; t = phenotypic effects of the imputed transcriptome; ε = residuals. Prediction accuracy improvement (i.e., y-axis) was derived by subtracting the prediction accuracy of the model y = g+ε from that of the model y = g+t+ε. The least squares line with 95% confidence band is based on a linear model that regressed prediction accuracy improvement on phenotypic variance explained by the imputed transcriptome. The p-value is for the t-test statistic (df=8) under the null hypothesis that the slope of the regression line is zero. **Panel b.** the large the estimated covariance between phenotypic effects of the genome and those of the imputed transcriptome (i.e., σ_gt_), the greater the gain in phenotypic prediction accuracy. Prediction accuracy was computed using the Pearson’s correlation coefficient between the observed and the predicted for two m odels of the form y = g+t+ε,but one assuming σ_gt_ = 0, i.e., GREML, and the other setting σ_gt_ as a free parameter for estimation, i.e., CORE GREML. Prediction accuracy improvement was derived by subtracting the prediction accuracy of GREML from that of CORE GREML. The least squares line with 95% confidence band is based on a linear model that regressed prediction accuracy improvement on σ_gt_ estimates. The p-value is for the t-test statistic (df=8) under the null hypothesis that the slope of the regression line is zero.

By comparing model fit by GREML and CORE GREML, we detected significant covariance between phenotypic effects of the regulatory regions and DHS for height and sitting height (Figure 3). Of note, the genomic partitioning model included three covariance terms, but two of these terms were not significant for height and sitting height, based on the Wald test with one degree of freedom. We therefore reduced the model for these two traits by dropping non-significant covariance terms, noting that the fit of the reduced model did not differ from the full model (*p* = 0.85 & 0.37 for height and sitting height, respectively). We subsequently used estimates from the reduced model for these two traits. In genome-transcriptome analyses, we found significant covariance between phenotypic effects of the genome and those of the imputed transcriptome for height, sitting height, heel bone mineral density, and diastolic blood pressure (Figure 4). We standardized all estimated covariance terms using respective variance estimates to derive correlation estimates (Figures 3 & 4 far right), noting that all significant estimates were positive and small to moderate in size, ranging from 0.14 (standing height in Figure 3) to 0.58 (heel bone mineral density in Figure 4). In a subsequent sensitivity analysis, all significant covariance terms emerging from the genomic partitioning and genome-transcriptome analyses remained after applying a rank-based inverse normal transformation to phenotypic observations (see p-values in Supplementary Figures 6 & 7). Thus, the estimated covariance terms were robust against the violation of the normality assumption held by GREML and CORE GREML.

**Figure 3.**
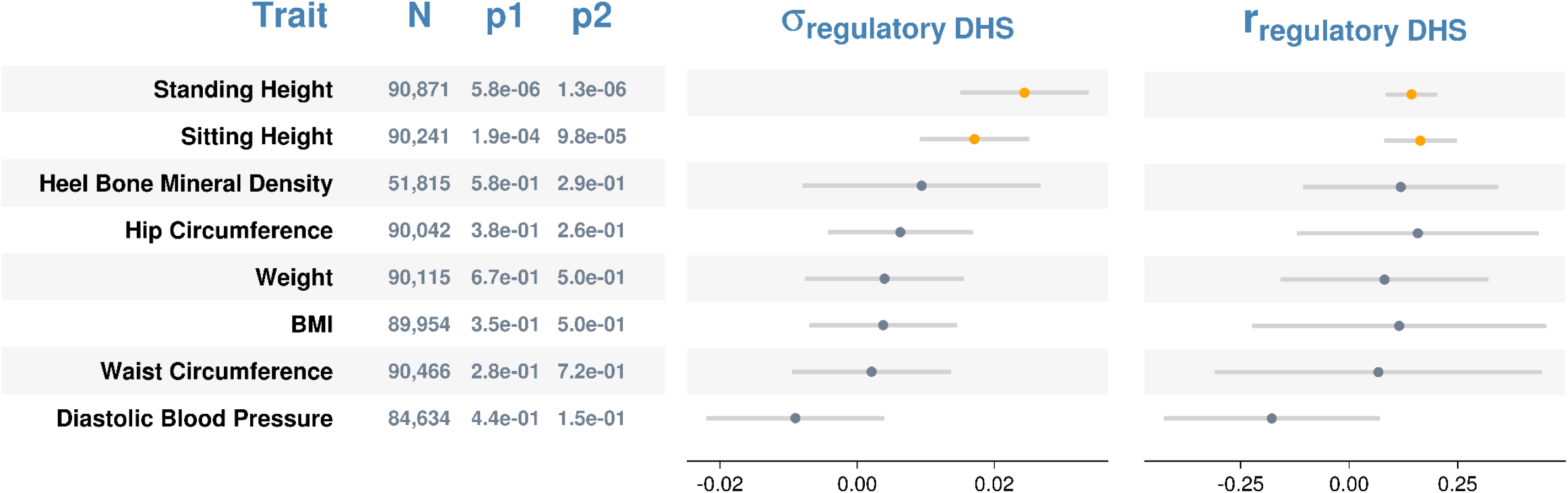
Estimated covariance (left) and correlations (right) between genetic effects attributable to the regulatory regions and those attributable to the DHS on phenotypes. Error bars are 95% confidence intervals. p1 = p-values from likelihood ratio tests that compare GREML with CORE GREML to detect σ_regulatory DHS_; p2 = p-values based on the Wald test statistic under the null hypothesis that r_regulatory DHS_= 0. Significant σ_regulatory DHS_ and r_regulatory DHS_ are highlighted in orange. Fluid intelligence and years of education are excluded because either the phenotypic effects of the regulatory regions or those of the DHS were not significant.

**Figure 4.**
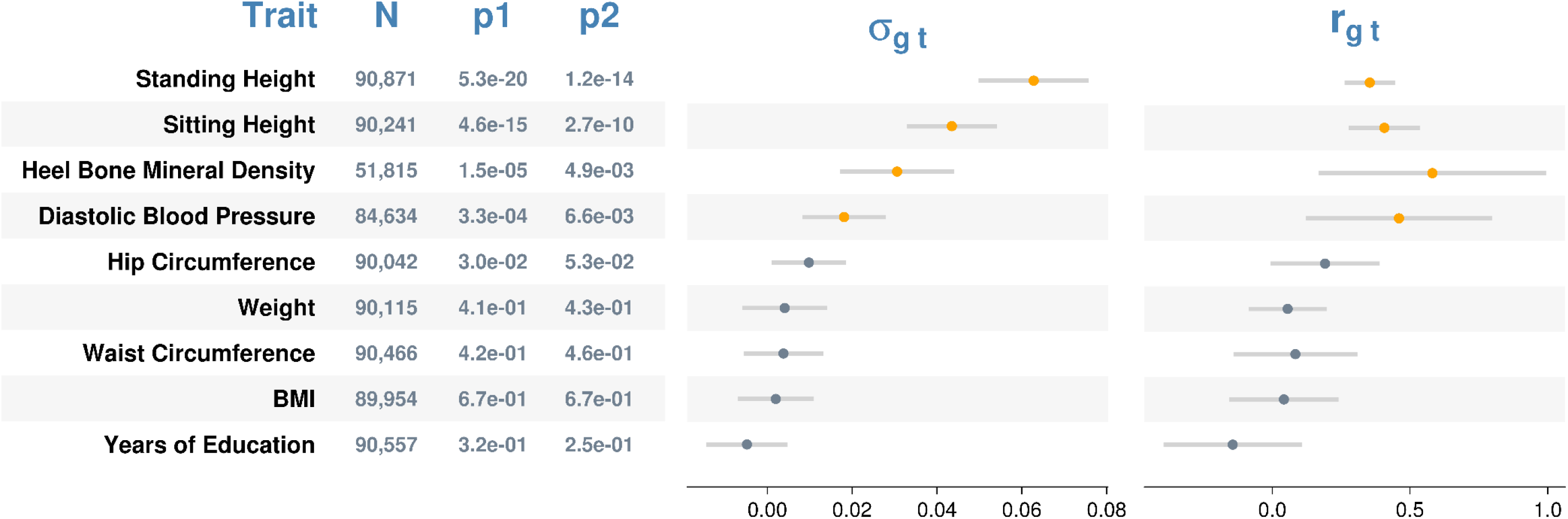
Estimated covariances (left) and correlations (right) between phenotypic effects of the genome and those of the imputed transcriptome. Error bars are 95% confidence intervals. p1 = p-values from likelihood ratio tests that compare GREML with CORE GREML to detect σ_gt_; p2 = p-values based on the Wald test statistic under the null hypothesis that r_gt_= 0. Significant σ_gt_ and r_gt_ are highlighted in orange. Fluid intelligence is excluded because the phenotypic effects of the imputed transcriptome on this trait was not significant after Bonferroni correction.

**Figure 5.**
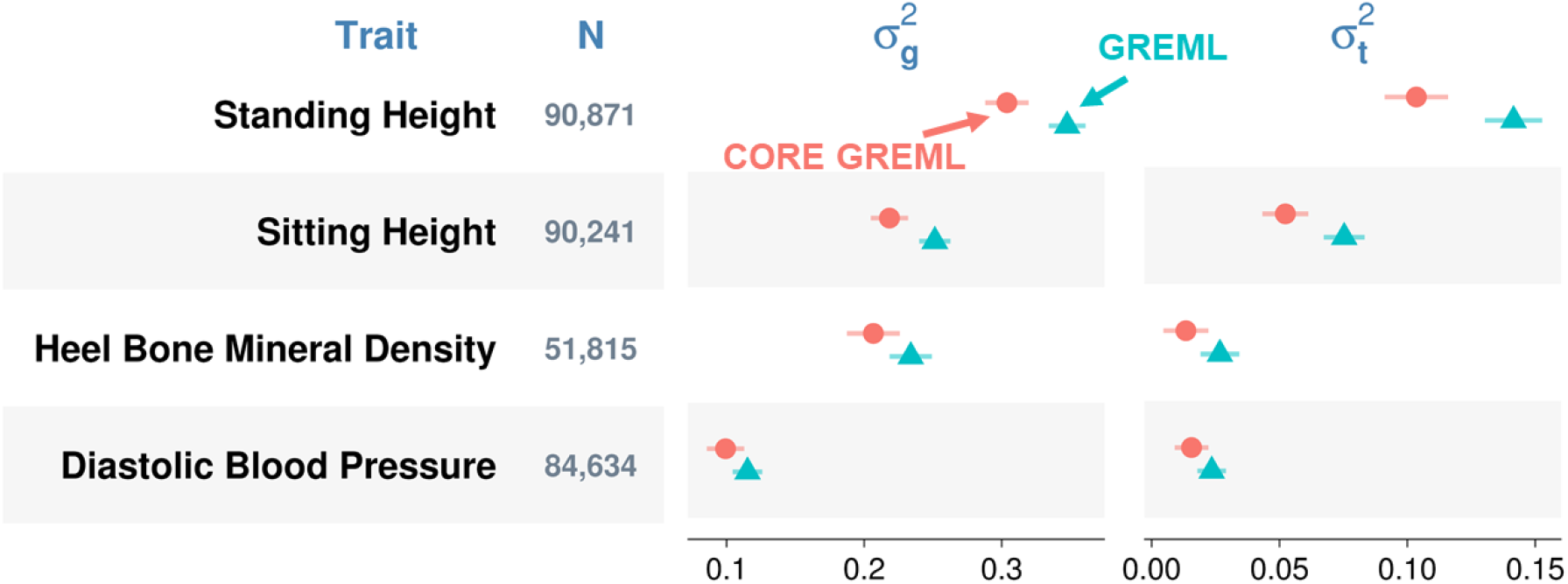
Variance component estimates by method from genome-transcriptome partitioning of phenotypic variance. 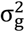 and 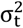 denote the phenotypic variances explained by the genome and by the imputed transcriptome, respectively. Error bars are 95% confidence intervals. Residual variance estimates are omitted for simplicity.

**Figure 6.**
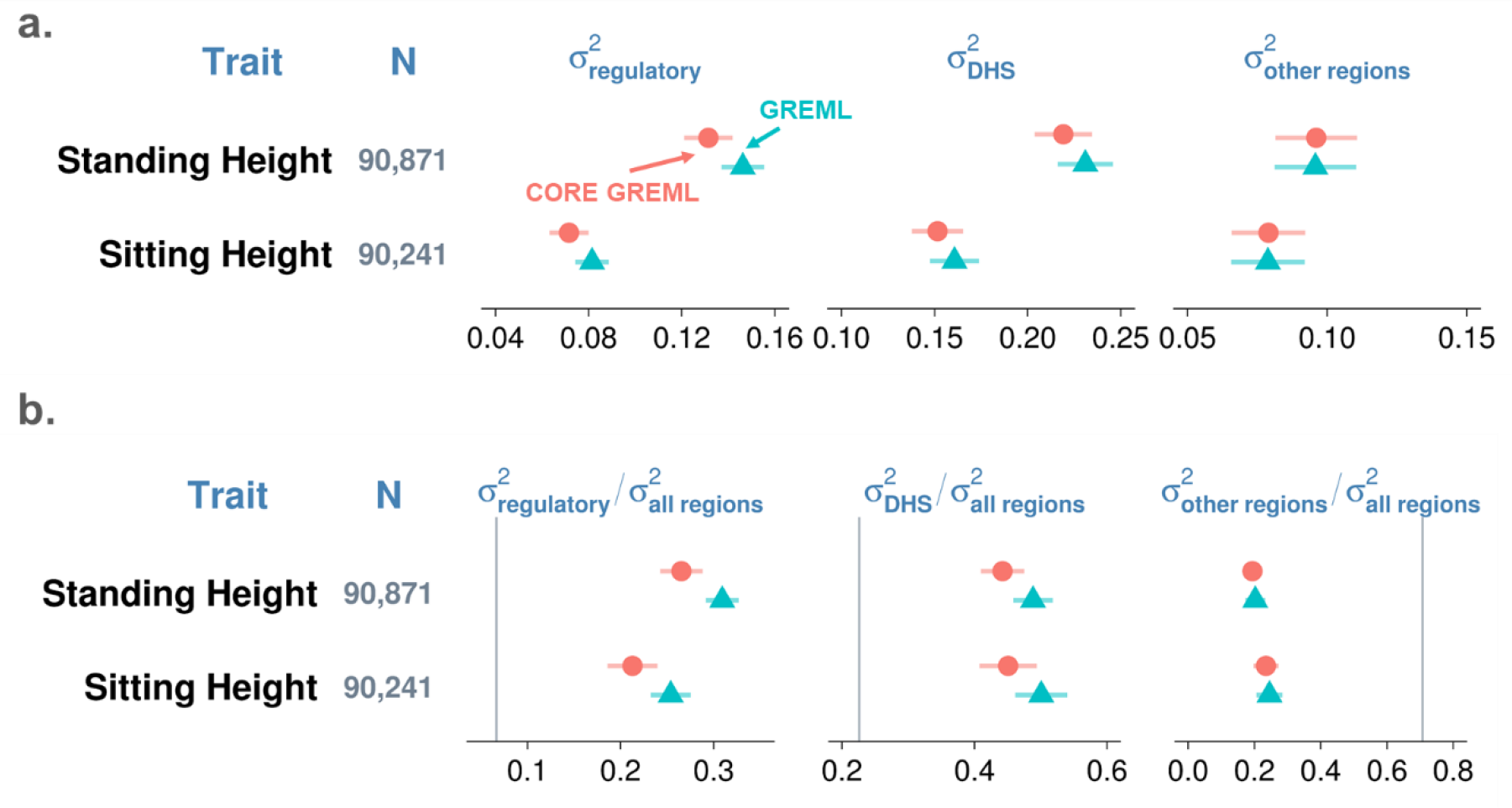
Model estimates by method from genomic partitioning analyses. Three functional regions of the genome are regulatory regions, DHS, and all other regions. 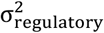, 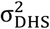, and 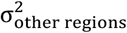 denote phenotypic variances explained by the three functional regions. a. variance component estimates; b. estimated proportions of total genetic variance attributable to three functional regions of the genome. Error bars are 95% confidence intervals. Vertical lines in panel b are percentages of SNPs from the three functional regions; conceptually, they are expected proportions of total genetic variance explained by the three functional regions of the genome assuming all genome-wide SNPs have an equal contribution to phenotypic variation. Residual variance estimates are omitted for simplicity.

To show that covariance terms estimated by CORE GREML are genuine biological parameters, we validated the covariance between phenotypic effects of the genome and the imputed transcriptome using the same 5-fold cross-validation procedure as before (i.e., for the validation of transcriptomic effects on phenotypes). In this instance, the phenotypic prediction accuracy of CORE GREML was compared against that of GREML. We chose genome-transcriptome analyses for validation since significant covariance emerged from four traits in contrast to two traits in genomic partitioning analyses. Figure 2 b. shows that the gain in the phenotypic prediction accuracy of CORE GREML relative to that of GREML grew as the magnitude of covariance estimates increased (*p* = 1.61e-05).

To show the impact of neglecting significant covariance terms, we compared variance component estimates by GREML with those by CORE GREML. In both genomic partitioning and genome-transcriptome analyses, variance estimates by GREML for correlated random effects were larger than those by CORE GREML (Figures 5 & 6a for significant results; see Supplementary Figures 6 & 7 for full results), noting that the differences in estimates between the two methods were proportional to the magnitude of covariance estimates. This is in line with our simulation results under positive covariance settings (cov > 0 in Supplementary Figures 2 & 3). As expected, neglecting covariance did not affect variance estimates for uncorrelated random effects. For example, variance estimates of the phenotypic effects of other genomic regions by CORE GREML for standing and sitting height agreed with those by GREML (Figure 6a). Similarly, for traits without any significant covariance term, there were minimal differences between GREML and CORE GREML estimates (Supplementary Figures 6 & 7), which aligned with simulation results under settings of zero covariance (cov = 0 in Supplementary Figures 2 & 3). Based on these observations, the larger variance estimates by GREML for correlated random effects relative to those by CORE GREML are most likely due to bias from neglecting the correlations between these random effects.

We also considered the impact of neglecting covariance between random effects on functions of variance component estimates, including 1) proportions of phenotypic variance explained by the genome (i.e., narrow-sense heritability) and by the imputed transcriptome (Supplementary Figure 8); 2) heritability partitioned by functional genomic region (Supplementary Figure 9); and 3) proportions of genetic variance by functional genomic region (Figure 6b for significant results; Supplementary Figure 10 for full results). These functions are useful for inferring the omic architecture of complex traits. For example, the relative phenotypic contributions of the genome and the imputed transcriptome can be inferred from the proportions of phenotypic variance explained by the two omes (note: these estimates are essentially identical to variance component estimates, because all traits have been standardized prior to analyses). The functional significance of SNPs from a given genomic region can be tested by assessing if the proportion of genetic variance attributable to the region is substantially higher than the the proportion of SNPs from the region^5^. Notably, GREML estimates of these functions were larger than CORE GREML estimates whenever there was a significant covariance term (Supplementary Figures 8-10); but the two methods agreed with each other otherwise. For example, the relative phenotypic contributions by the genome and the imputed transcriptome inferred from GREML were larger than those inferred from CORE GREML for heel bone mineral density, diastolic blood pressure, sitting height and standing height (Figure 8). Similarly, the functional significance of SNPs from regulatory regions inferred by GREML was larger than that by CORE GREML for sitting height and standing height (Figure 6b). Taken together, our results indicate that GREML can lead to incorrect inferences on the underlying architecture of complex traits unless correlations between random effects are properly modelled.

It is important to note that kernel matrices used for variance-components estimation in a LMM can be similar, for instance, due to linkage disequilibrium in a genomic partitioning analysis. In fact, the correlations between off-diagonal entries of kernel matrices used in our analyses are moderate to high (0.35-0.98; see Supplementary Table 6). This similarity might give rise to the covariance between random effects. However, this possibility is unlikely for at least two reasons. First, if covariance is driven by the similarity between kernel matrices, then we would expect that in the null setting of our simulations, type I error rate is inflated, given the high similarities between kernel matrices in our analyses. Contrary to this, we found that type I error rate is controlled. Second, the kernel matrix constructed using genotypes from DHS is more similar to the kernel matrix constructed using genotypes from other regions than to the one for regulatory regions; but significant covariance was only detected between effects of regulatory regions and those of DHS on standing height and sitting height. Therefore, covariance between random effects is unlikely driven by the similarity between kernel matrices for variance-components estimation.

## Discussion

When applying the classic LMMs for standard heritability estimation, where phenotypic variance is only partitioned into genetic and residual variances, the model assumption of negligible covariance between random effects (i.e., genetic and residual effects) may be met in many cases. However, when phenotypic variance is further partitioned, for example, by functional genomic region or omic layer, using a model with multiple random effects, the covariance terms between these random effects can be substantial, as what we demonstrated using the genomic partitioning analyses and the genome-transcriptome partitioning analyses for complex traits. Unless these non-negligible covariance terms are properly accounted for, variance-components estimation would be biased, resulting in misleading inferences on the latent omic architecture of complex traits, as shown by simulation results. Therefore, we recommend that covariance terms between random effects need to be carefully checked and properly modelled for genomic analyses of complex traits.

CORE GREML can serve as a useful tool for detecting and estimating covariance terms between random effects, as demonstrated using analyses of both simulated and real data. Prior to the proposal of CORE GREML, there have been several attempts to relax the assumption of independence between random effects^20,21^, but they are specific to experimental studies and are not readily applicable to genome-wide analyses for human complex traits. To our knowledge, CORE GREML is the first of its kind in variance partitioning analyses, which correctly models the covariance between random effects.

We demonstrated the use of CORE GREML in genomic partitioning analyses and genome-transcriptome partitioning of phenotypic variance, and found that significant covariance terms mostly emerged from the latter (for 4 traits out of 10). While genome-transcriptome associations have been reported by numerous studies^12–15^, they were based on a limited number of SNPs and genes. In contrast, the association estimates by CORE CREML were based on aggregated effects of genome-wide SNPs and those of all available gene expression levels jointly on phenotypes, thereby providing an overall picture of the proportion of phenotypic variance shared by the whole genome and the transcriptome.

Of note, our study used imputed, as opposed to measured, gene expression. As demonstrated by our cross-validation results, the imputed transcriptome already improves phenotypic prediction accuracy, hence it allows phenotype forecasting, for example, for newborns, solely on the basis of genotype information. However, our analyses using the imputed gene expression would only have captured part of transcriptomic effects on phenotypes; hence the phenotypic variance explained by the transcriptome would have been underestimated. When actual gene expression levels become available for future analyses, we expect an additional gain in explained phenotypic variance. The use of the imputed transcriptome may also explain that although the model with an additional random term for phenotypic effects of the imputed transcriptome fit the data much better than the model without, the two models explained the same amount of total phenotypic variance. Notably, this result aligns with a recent notion of total genetic effects on complex traits, which is partitioned into genetic effects mediated by gene expression and ones not^22^. While the former is essentially the effects of imputed gene expression on phenotypes, the latter is the effects of common SNPs (see Equation 3 in Yao et al.^22^). Despite the conceptual similarity, our study is different from Yao et al. in two key aspects. First, implied from the model in Yao et al., gene expression levels were based on SNPs at cis-eQTLs only; but in our study gene expression levels were computed using genome-wide SNPs. This may explain that the estimated phenotypic variance due to imputed gene expression in their study tends to be smaller than that in ours for BMI, standing height, heel bone mineral density and years of education. Second, unlike CORE GREML used in our study, the model used by Yao et al. does not account for covariance between effects of the genome and the imputed transcriptome on phenotypes.

Importantly, the proposed CORE GREML can be used to analyse and dissect the shared effects among omic layers, beyond the genome and the transcriptome, including proteome, metabolome and exposome, when multi-omics data become available. We anticipate, based on our genome-transcriptome analyses, that covariance between phenotypic effects of omic layers is a key parameter, such that CORE GREML will be an important tool for multi-omics partitioning analyses. Other potential applications of CORE GREML include phenotypic variance partitioning by chromosome^23^ or minor allele frequency bin^24,25^, where correlations between phenotypic effects of random terms are intuitive. Even for the simplest partitioning of phenotypic variance that separates genetic variance apart from residual variance, CORE GREML can be useful, if genetic effects and residuals are correlated due to confounding, associations or interactions between genetic and environmental effects^26^. However, CORE GREML is computationally intensive, such that estimation becomes infeasible for models including a large number of random effects (e.g., the baseline model in Finucane et al.^8^ with >100 random-effects terms). Further studies are required to develop computationally efficient algorithms for CORE GREML, e.g. using summary statistics. In addition, we only validated CORE GREML for quantitative traits in this study. Validation of our method for binary traits is required in future studies.

Finally, we showed, using simulations, that misspecification of LD and MAF dependent genetic architecture can cause substantial bias in variance-components estimation by CORE GREML, although covariance estimation seems robust. However, the likelihood of the true estimation model in general is much greater than a wrong model, suggesting that likelihood-based comparisons of models that assume different genetic architectures is a useful way to reduce the chance of mis-specifying genetic architecture. For demonstration, we fitted the GCTA model and the LDAK model with the recommended default setting^19^ and found the GCTA model in general had a better fit than the LDAK model for our traits. In the absence of the knowledge of the true genetic architecture, however, it is recommended to more systematically vary parameter settings of the LDAK model (as in^19^) and choose the best fitting model via likelihood comparison before applying CORE GREML.

In this study, we introduce a generalised GREML, referred to as CORE GREML, that relaxes the assumption of independence between phenotypic effects of random terms held by classic mixed-effects models for variance component analyses. Using both simulations and real data, we showed that in the presence of non-negligible covariance terms, CORE GREML improved genomic partitioning and multi-omics partitioning analyses by the conventional GREML. We conclude that the covariance between random effects for analysis of complex traits is a key parameter for estimation, and hence, recommend that covariance terms should be carefully checked and properly modelled.

## Methods

### Generalizing variance-covariance matrix of phenotypic observations

A LMM can be written as

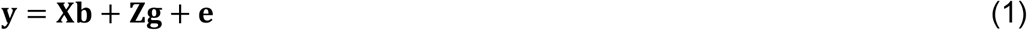

where **y** is a vector of trait phenotypes, **b** is a vector of fixed effects, **g** is a vector of additive genetic effects and **e** is a vector of residual effects. **X** and **Z** are incidence matrices. The random effects, **g** and **e**, are assumed to be normally distributed with mean zeros and variances 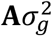 and 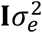, respectively, where **A** and **I** are the genetic relationship kernel matrix^2,27,28^ and an identity matrix, respectively. The variance-covariance matrix of all observations, var(**y**), can be written as

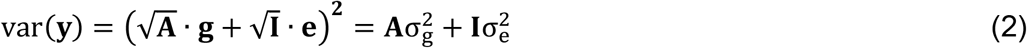

where 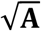 is the Cholesky decomposition of the genetic relationship kernel matrix with 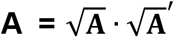 and 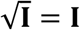 by the identity matrix property. This is the standard definition of variance covariance matrix used in LMM, which assumes no correlation between **g** and **e**, i.e. cor(**g**, **e**) = 0.

Relaxing this classical assumption, eq. (2) can be expressed as

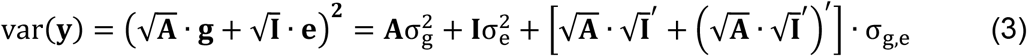

where *σ*_*g,e*_ is the covariance between **g** and **e**, i.e. 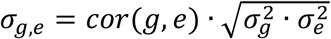.

When considering multiple random effects in the LMM (e.g. genomic partitioning approach), the model can be written as

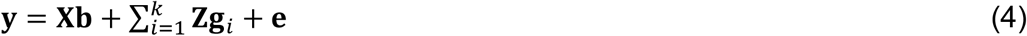

where **g**_*i*_ is the random genetic effects of the *i*th pre-defined functional category, e.g. regulatory regions.

Such mixed-effects models with multiple random terms typically assume no correlation between **g**_*i*_ and **g**_*j*_. However, this assumption can be violated if effects of two categories on phenotypes are associated, for example, through the same gene pathway. We relax this assumption and write the variance-covariance matrix of all observations, var(**y**), as

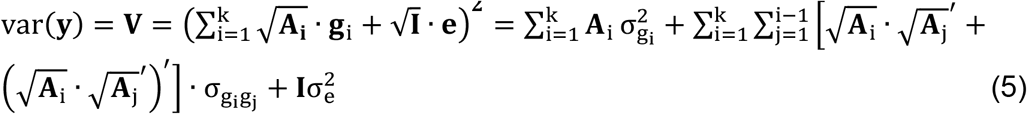

where **A**_*i*_ is the genetic relationship kernel matrix constructed using SNPs from the *i*th functional category, 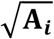 is the Cholesky decomposition of **A**_*i*_ and 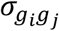 is the genetic covariance between **g**_*i*_ and **g**_*j*_. It is noted that the correlation term between genetic effects (**g**_*i*_) and residuals (**e**) is not included and hence assumed to be zero in Eq. (4), which is usually valid, although it is possible to parameterise this term.

The log likelihood of the proposed model, which can be generally applied to Eq. (1) and (4), is

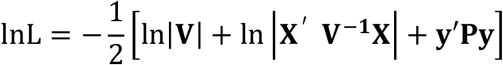

where *ln* is the natural log and | | the determinant of the associated matrices. The projection matrix is defined as **P** = **V**^−1^ − **V**^−1^**X**(**X**^′^**V**^−1^**X**)^−1^**X**^′^**V**^−1^. By maximising the log likelihood, the direct average information algorithm^29,30^ can be used to obtain CORE GREML estimates of parameters including the covariance terms between random effects.

This CORE GREML approach can be easily extended to phenotypic variance partitioning using multi-omics data, e.g. genome-transcriptome analyses (see Genome-Transcriptome Partitioning Model section below).

### Heritability

For Eq. (1), the standard definition of heritability is

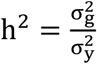

where the phenotypic variance is 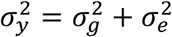 in the absence of cor(**g**, **e**). When there is non-negligible cor(**g**, **e**), the phenotypic variance should be written as 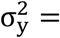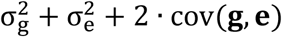.

For Eq. (4), a general expression of heritability for the *i*th genetic component is

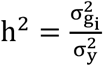

where 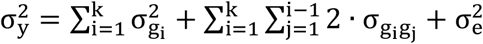.

Using the Delta method (see^31^), the sampling variance of heritability for this example can be obtained as

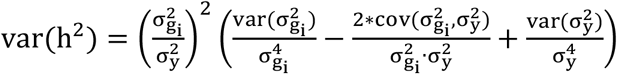

where 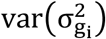, 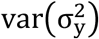 and 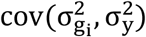 can be obtained from the average information matrix^32^ of CORE GREML.

### Correlation between two random effects

The correlation between two random (genetic) effects 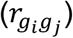 can be defined as the genetic covariance scaled by the square root of the product of the genetic variances of the two random effects, that is

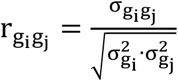

Using the Delta method (see^31^), the sampling variance of genetic correlation can be obtained as

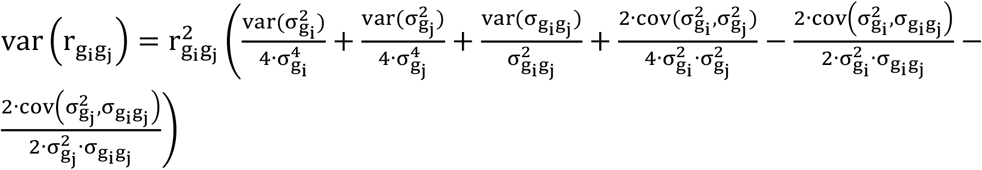

where the variance and covariance terms used are from the information matrix of CORE GREML.

### Real data analysis

#### Genotype data

The UK biobank (project approval number 14575) contains health-related data from ~ 500,000 participants aged between 40 and 69, who were recruited throughout the UK between 2006 and 2010^33^. Prior to data analysis, we applied stringent quality control to exclude unreliable genotypic data. We filtered SNPs with an INFO score < 0.6, a MAF < 0.01, a Hardy-Weinberg equilibrium p-value <1e-4, or a call rate < 0.95. We then selected HapMap3 SNPs, which are known to yield reliable and robust estimates of SNP-based heritability^25,34,35^, for downstream analyses. We filtered individuals who had a genotype-missing rate > 0.05, were non-white British ancestry, or had the first or second ancestry principal components outside six standard deviations of the population mean. We also applied relatedness-cut-off quality control to exclude one of any pair of individuals with a genomic relationship > 0.025. From the remaining individuals, we selected those who were included in both the first and second release of UK biobank genotype data. We calculated the discordance rate of imputed genotypes between the two versions and excluded individuals with a discordance rate > 0.05. Eventually, genotypes of 1,131,002 SNPs from 91,472 individuals remained for data analysis.

#### Phenotype data

To preclude negligible heritability as a possibility for negative findings (i.e., no covariance between random effects), we deliberately chose ten UK biobank traits available to us with the largest heritability estimates by an independent open source (https://nealelab.github.io/UKBB_ldsc/), which included standing height, sitting height, body mass index (BMI), heel bone mineral density, fluid intelligence, weight, waist circumference, hip circumference, diastolic blood pressure and years of education^36^. Heritability estimates for all selected traits were at least 20 times greater than their standard errors to ensure that they were significantly different from zero. We further verified SNP-based heritability of these traits using GREML and estimates are shown in Supplementary Figure 1.

Prior to model fitting, phenotypic data were prepared in three sequential steps: 1) adjustment for age, sex, birth year, social economic status (by Townsend Deprivation Index), population structure (by the first ten principal components of the estimated genomic relationship matrix), assessment centre, and genotype batch using linear regression; 2) standardization; and 3) removal of data points outside +/− 3 standard deviations from the mean. The distributions of phenotypes of the ten traits are shown in Supplementary Figure 11. We noted mild to strong deviations from normality for traits such as BMI and years of education. This motivated a subsequent sensitivity analysis to test the robustness of our findings against the violation of the normality assumption held by GREML and CORE GREML. Specifically, we applied a rank-based inverse normal transformation to phenotypes of all traits and repeated our analyses on the transformed phenotypes.

#### Functional annotation of the genome

The genome was annotated using three pre-defined functional categories (https://data.broadinstitute.org/alkesgroup/ANNOTATIONS): 1) regulatory regions that consist of coding regions, untranslated regions and promotors; 2) DNase I hypersensitivity sites (DHS); and 3) all other regions. We assigned each SNP into one of the three categories, resulting in 75,396 SNPs in the regulatory regions, 255,665 in the DHS and 799,935 in all other regions. Prior to the assignment, genotype data were quality-controlled (see above for details).

#### Gene expression imputation

Using PrediXcan^18^, we imputed expression levels of 2,028 to 9,630 genes for 43 non-sex-specific tissues (Supplementary Table 2) by projecting estimated SNP effects of expression onto genotypes of 2,392,786 SNPs for 91,472 individuals from the UK Biobank. Selected SNPs had an INFO score > 0.6, a minor allele frequency > 0.01, a p-value for the Hardy-Weinberg test < 0.0001, and missingness < 0.05. SNP effect estimates were sourced from GTEx v7 models (2018-01-08 release; http://predictdb.org/) that were trained using 2,496,846 SNPs of European individuals from the Genotype-Tissue Expression project^37^.

#### Variance partitioning models

The phenotypic variance of each selected trait was partitioned using two separate random-effects models (see below for model description). The ‘genome-transcriptome’ model partitions phenotypic variance into variation from the genome, the transcriptome and unknown sources (i.e., residual variance), whereas the ‘genomic partitioning’ model assumes phenotypic variation comes from the genome and residuals and further partitions genetic variance by functional category of the genome. Three functional categories were under consideration, namely, regulatory regions (encompassing coding regions, untranslated regions and promotors), DNase I hypersensitivity sites (DHS) and all other regions.

Each partitioning model was fitted using the conventional method, i.e., GREML, and the proposed alternative, i.e., CORE GREML. Essentially, GREML sets all covariance terms between random effects to zero, whereas CORE GREML treats these terms as free parameters for estimation. To detect significant covariance terms, we performed likelihood ratio tests to determine whether the model fit by CORE GREML was better than that by GREML.

### a. Genome-Transcriptome Partitioning Model

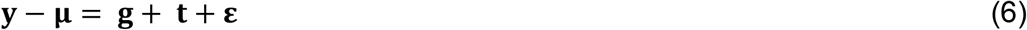

Where **y** is a n × 1 vector of phenotype data and their deviations from the grand mean, **μ**, are decomposed into three vectors consisting of phenotypic effects of the genome, 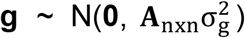, those of the transcriptome, 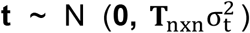, and residuals, 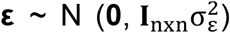. The terms, 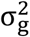, 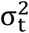 and 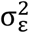 denote phenotypic variances attributable to the genome, the transcriptome and residuals, respectively. **A**_nxn_ and **T**_nxn_ are relationship kernel matrices and **T**_nxn_ is an identity matrix. **A**_nxn_ is derived by 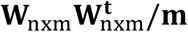, and **T**_nxn_ by 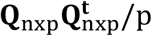, where **W**_nxm_ contains standardized genotype information of *m* (=1,131,002) SNPs for *n* (= 91,472) individuals, and **Q**_nxp_ contains standardized imputed expression of *p* (=227,664) genes collapsed across 43 tissues for the *n* individuals. Essentially, entries of **A**_nxn_ and *T*_nxn_ describe pairwise similarities between individuals based on their genotypes and imputed gene expression, respectively.

The variance covariance matrix of phenotypic observations is

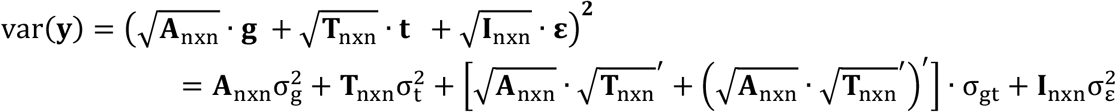

where σ_gt_ is the covariance between the effects of the genome and of the transcriptome on phenotypes. Here we assume no correlation between residuals and genomic or transcriptomic effects.

### b. Genomic Partitioning Model

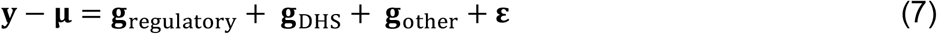

where **y** is a n × 1 vector of phenotype data, and their deviations from the grand mean, **μ**, are decomposed into genetic effects due to regulatory, 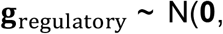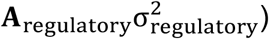, those due to DHS, 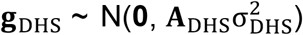, those due to other genomic regions 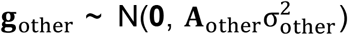, and residuals, 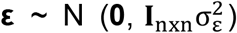. The terms, 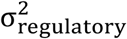, 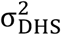, 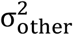 and 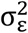 denote phenotypic variances attributable to the three functional regions and residuals, respectively. The kernel matrices, **A**_regulatory_, **A**_DHS_ and **A**_other_ were constructed using 75,396 SNPs from regulatory regions, 255,665 from DHS, and 799,935 from all other genomic regions. **I** is a n × n identity matrix.

The variance covariance matrix of phenotypic observations is

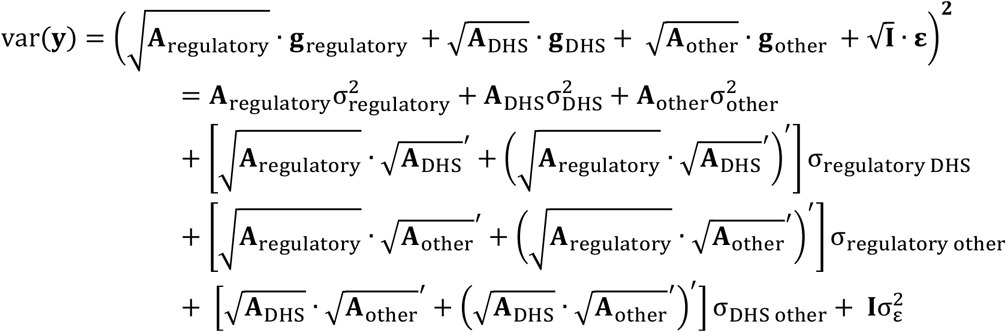

where σ_ij_ is the covariance between genetic effects of functional regions *i* and *j*, for *i* & *j* ϵ {regulatory regions, DHS, other regions} and *i* ≠ *j*. We assume no correlation between residuals and any of the genetic effects.

### Simulation

To validate CORE GREML, we simulated 500 replicates of phenotypic data using the two variance partitioning models shown above under each of three parameter settings: zero (i.e., null setting), positive and negative covariance between phenotypic effects of random terms in the variance partitioning model. Simulations were based on quality-controlled genotype data and imputed transcriptome data from a random sample of 10,000 UK biobank individuals. The genotype data contained a total of 1,131,002 SNPs (see *genotype data* above), and the imputed transcriptome contained imputed expressions of 227,664 genes collapsed cross 43 non-sex specific tissues (see *gene expression imputation* above).

For genome-transcriptome analysis, phenotypes were simulated using eq. 6 according to the following variance-covariance structure of random effects:

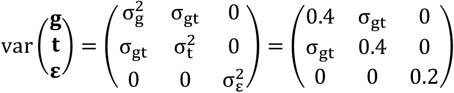

where the value of σ_gt_ varied across parameter settings (Supplementary Table 1).

For genomic partitioning analysis, phenotypes were simulated using eq. 6 according to the following variance-covariance structure of random effects:

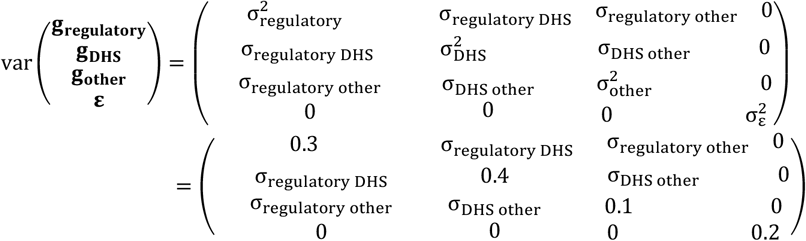

where the values of covariance terms varied across parameter settings (Supplementary Table 1).

For each replicate, we fitted the two variance partitioning models using both GREML and CORE GREML as for analysis of real data. Under the null setting, we assessed if CORE GREML can detect covariance term(s) at a controlled rate of type I errors, by comparing the model fit of CORE GREML with that of GREML using likelihood ratio tests. Under all settings, we assessed if CORE GREML can produce unbiased estimates of variance and covariance components. To show the impact of neglecting genuine covariance terms, we also provided parameter estimates by GREML.

To test the sensitivity of CORE GREML estimation to a wrong assumption about the genetic architecture in the estimation, we also simulated phenotypes under genetic architecture. Following past studies ^19,38^, we parameterized genetic architecture by minor allele frequency and linkage disequilibrium, assuming the variance of SNP-specific effects on phenotypes, var(*β*), for any given SNP *i*, is proportional to its linkage disequilibrium score, *w*, and minor allele frequency, *f*, expressed as 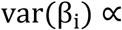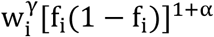, where α and γ control the extents to which *w* and *f* affect var(*β*). By altering values of *α* (either −1 or −0.25) and *γ* (either 0 or 1), we simulated phenotypic data under three different genetic architectures (*α* = −1, *γ* = 1; *α* = −0.25, *γ* = 1; *α* = −0.25, *γ* = 0), each including three scenarios of the covariance between phenotypic effects of the genome and those of the transcriptome, namely, null (σ_gt_ = 0), positive (σ_gt_ = 0.2) and negative ( σ_gt_ = −0.2), as for the CORE GREML validation simulations (Supplementary Table 1; note the genetic architecture for the CORE GREML validation simulations is under the setting *α* = −1, *γ* = 0). Each scenario had 500 replicates of simulated phenotypic data (each with n = 10,000). For a given genetic architecture, we fitted two estimation models, both using CORE GREML, but one assuming values of *α* and *γ* the same as those of the simulation model (i.e., ‘true model’) and the other always assuming *α* = −1, *γ* = 0 (i.e., ‘wrong model’).

## Acknowledgements

This research is supported by the Australian Research Council (DP190100766, FT160100229) and the Australian National Health and Medical Research Council (1087889). The authors would like to thank staff and participants of the UK Biobank for their important contributions. The UK biobank was approved by the NHS National Research Ethics Service North West (11/NW/0382), and our access was under the reference number 14575. Work was performed using computational resources provided by the Australian Government through Raijin under the National Computational Merit Allocation Scheme (NCMAS).

## URLs

UK Biobank: http://www.ukbiobank.ac.uk/

MTG2 for model fitting: https://sites.google.com/site/honglee0707/mtg2

PrediXcan for transcriptome imputation: https://github.com/hakyimlab/PrediXcan

Estimated SNP effects on gene expression from GTEx v7 models: http://predictdb.org/

## Code availability

Example code along with related files for fitting CORE GREML can be found from the following site: https://sites.google.com/site/honglee0707/mtg2

## Reference

1. Manolio, T.A. et al. Finding the missing heritability of complex diseases. Nature 461, 747 (2009).

2. Yang, J. et al. Common SNPs explain a large proportion of the heritability for human height. Nature genetics 42, 565 (2010).

3. Henderson, C.R., Kempthorne, O., Searle, S.R. & Von Krosigk, C. The estimation of environmental and genetic trends from records subject to culling. Biometrics 15, 192–218 (1959).

4. Fisher, R.A. XV.—The correlation between relatives on the supposition of Mendelian inheritance. Earth and Environmental Science Transactions of the Royal Society of Edinburgh 52, 399–433 (1919).

5. Gusev, A. et al. Partitioning heritability of regulatory and cell-type-specific variants across 11 common diseases. The American Journal of Human Genetics 95, 535–552 (2014).

6. Hasin, Y., Seldin, M. & Lusis, A. Multi-omics approaches to disease. Genome biology 18, 83 (2017).

7. Bulik-Sullivan, B.K. et al. LD Score regression distinguishes confounding from polygenicity in genome-wide association studies. Nature genetics 47, 291 (2015).

8. Finucane, H.K. et al. Partitioning heritability by functional annotation using genome-wide association summary statistics. Nature genetics 47, 1228 (2015).

9. Johannesson, M., Magnusson, P., Ikram, M. & Visscher, P. Equivalence of LD-Score Regression and Individual-Level-Data Methods. Behavioural Genetics 47, 642–719 (2017).

10. Ni, G. et al. Estimation of genetic correlation via linkage disequilibrium score regression and genomic restricted maximum likelihood. The American Journal of Human Genetics 102, 1185–1194 (2018).

11. Bulik-Sullivan, B. et al. An atlas of genetic correlations across human diseases and traits. Nature genetics 47, 1236 (2015).

12. Mercer, T.R. et al. DNase I–hypersensitive exons colocalize with promoters and distal regulatory elements. Nature genetics 45, 852 (2013).

13. Cooper, S.J., Trinklein, N.D., Anton, E.D., Nguyen, L. & Myers, R.M. Comprehensive analysis of transcriptional promoter structure and function in 1% of the human genome. Genome research 16, 1–10 (2006).

14. Shu, W., Chen, H., Bo, X. & Wang, S. Genome-wide analysis of the relationships between DNaseI HS, histone modifications and gene expression reveals distinct modes of chromatin domains. Nucleic acids research 39, 7428–7443 (2011).

15. Wang, Y.-M. et al. Correlation between DNase I hypersensitive site distribution and gene expression in HeLa S3 cells. PloS one 7, e42414 (2012).

16. Giambartolomei, C. et al. Bayesian test for colocalisation between pairs of genetic association studies using summary statistics. PLoS genetics 10, e1004383 (2014).

17. Farh, K.K.-H. et al. Genetic and epigenetic fine mapping of causal autoimmune disease variants. Nature 518, 337 (2015).

18. Gamazon, E.R. et al. A gene-based association method for mapping traits using reference transcriptome data. Nature genetics 47, 1091 (2015).

19. Speed, D. et al. Reevaluation of SNP heritability in complex human traits. Nature genetics 49, 986 (2017).

20. Dumont, C., Chenel, M. & Mentré, F. Influence of covariance between random effects in design for nonlinear mixed-effect models with an illustration in pediatric pharmacokinetics. Journal of biopharmaceutical statistics 24, 471–492 (2014).

21. Frossard, J. & Renaud, O. The correlation structure of mixed effects models with crossed random effects in controlled experiments. arXiv preprint arXiv:1903.10766 (2019).

22. Yao, D.W.,O’connor, L.J., Price, A.L. & Gusev, A. Quantifying genetic effects on disease mediated by assayed gene expression levels. BioRxiv, 730549 (2019).

23. Yang, J. et al. Genome partitioning of genetic variation for complex traits using common SNPs. Nature genetics 43, 519 (2011).

24. Yang, J. et al. Genetic variance estimation with imputed variants finds negligible missing heritability for human height and body mass index. Nature genetics 47, 1114 (2015).

25. Lee, S.H. et al. Estimation of SNP heritability from dense genotype data. The American Journal of Human Genetics 93, 1151–1155 (2013).

26. VanderWeele, T.J., Ko, Y.-A. & Mukherjee, B. Environmental confounding in gene-environment interaction studies. American journal of epidemiology 178, 144–152 (2013).

27. Amin, N., Van Duijn, C.M. & Aulchenko, Y.S. A genomic background based method for association analysis in related individuals. PloS one 2, e1274 (2007).

28. VanRaden, P.M. Efficient methods to compute genomic predictions. Journal of dairy science 91, 4414–4423 (2008).

29. Lee, S.H. & Van Der Werf, J.H. An efficient variance component approach implementing an average information REML suitable for combined LD and linkage mapping with a general complex pedigree. Genetics Selection Evolution 38, 25 (2006).

30. Yang, J., Lee, S.H., Goddard, M.E. & Visscher, P.M. GCTA: a tool for genome-wide complex trait analysis. The American Journal of Human Genetics 88, 76–82 (2011).

31. Lynch, M. & Walsh, B. Genetics and analysis of quantitative traits, (Sinauer Sunderland, MA, 1998).

32. Gilmour, A.R., Thompson, R. & Cullis, B.R. Average information REML: an efficient algorithm for variance parameter estimation in linear mixed models. Biometrics, 1440–1450 (1995).

33. Sudlow, C. et al. UK biobank: an open access resource for identifying the causes of a wide range of complex diseases of middle and old age. PLoS medicine 12, e1001779 (2015).

34. Lee, S.H. et al. Genetic relationship between five psychiatric disorders estimated from genome-wide SNPs. Nature genetics 45, 984 (2013).

35. Ripke, S. et al. Genome-wide association analysis identifies 13 new risk loci for schizophrenia. Nature genetics 45, 1150 (2013).

36. Okbay, A. et al. Genome-wide association study identifies 74 loci associated with educational attainment. Nature 533, 539 (2016).

37. Lonsdale, J. et al. The genotype-tissue expression (GTEx) project. Nature genetics 45, 580 (2013).

38. Hou, K. et al. Accurate estimation of SNP-heritability from biobank-scale data irrespective of genetic architecture. Nat Genet 51, 1244–1251 (2019).

