## Supplementary Materials for "CORE GREML: Estimating covariance between random effects in linear mixed models for genomic analyses of complex traits"

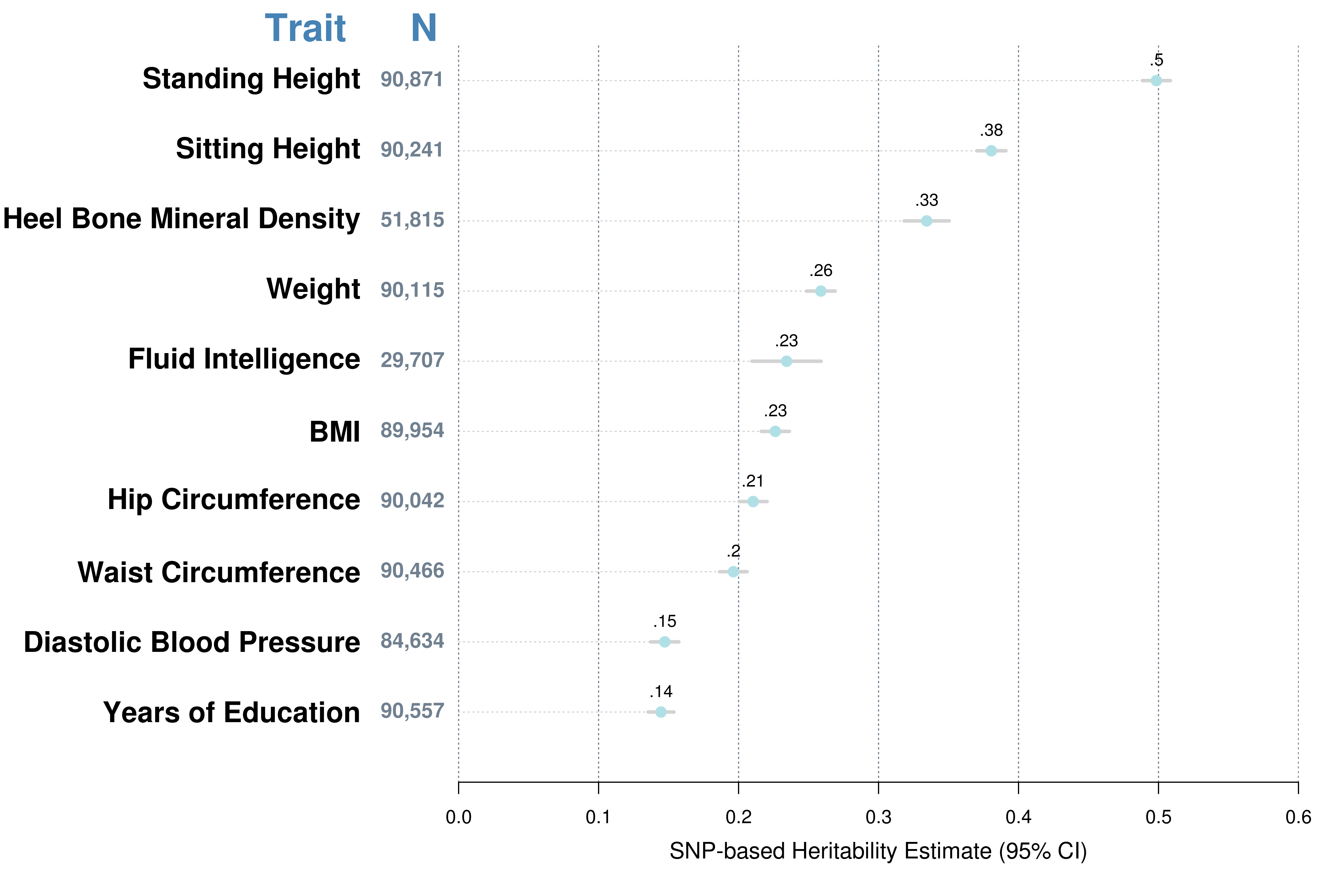
Supplementary Figure 1. SNP-based heritability estimates of ten selected traits from the UK Biobank.

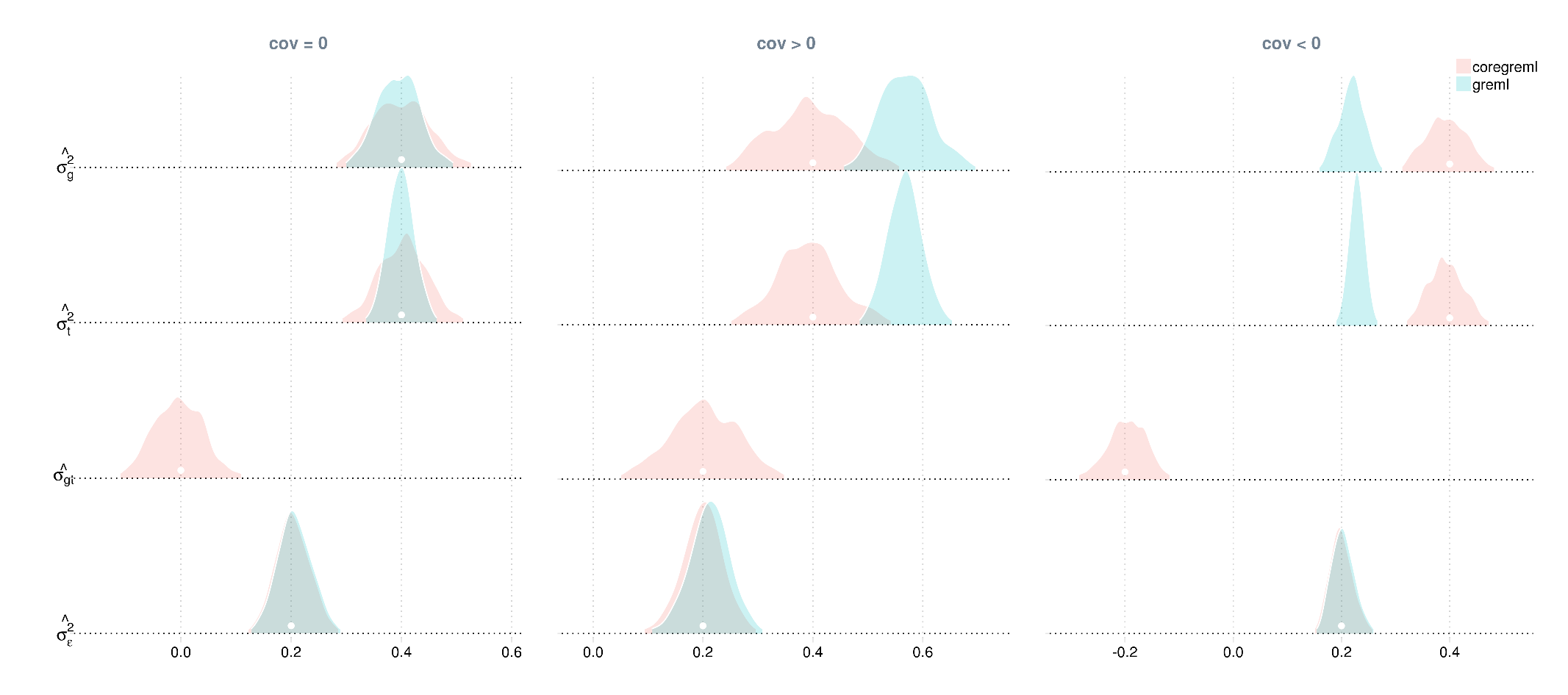

Supplementary Figure 2. Sampling distributions of model parameters by estimation method for genome-transcriptome partitioning of phenotypic variance. Five-hundred replicates of phenotypic data (n=10,000) were simulated under each of three parameter settings, where the covariance between phenotypic effects of the genome and the imputed transcriptome was zero (cov=0), positive (cov>0) and negative (cov<0). For each replicate, model parameters were estimated by the traditional method, i.e., GREML, and the proposed method, i.e., CORE GREML. True values of model parameters are in dots.

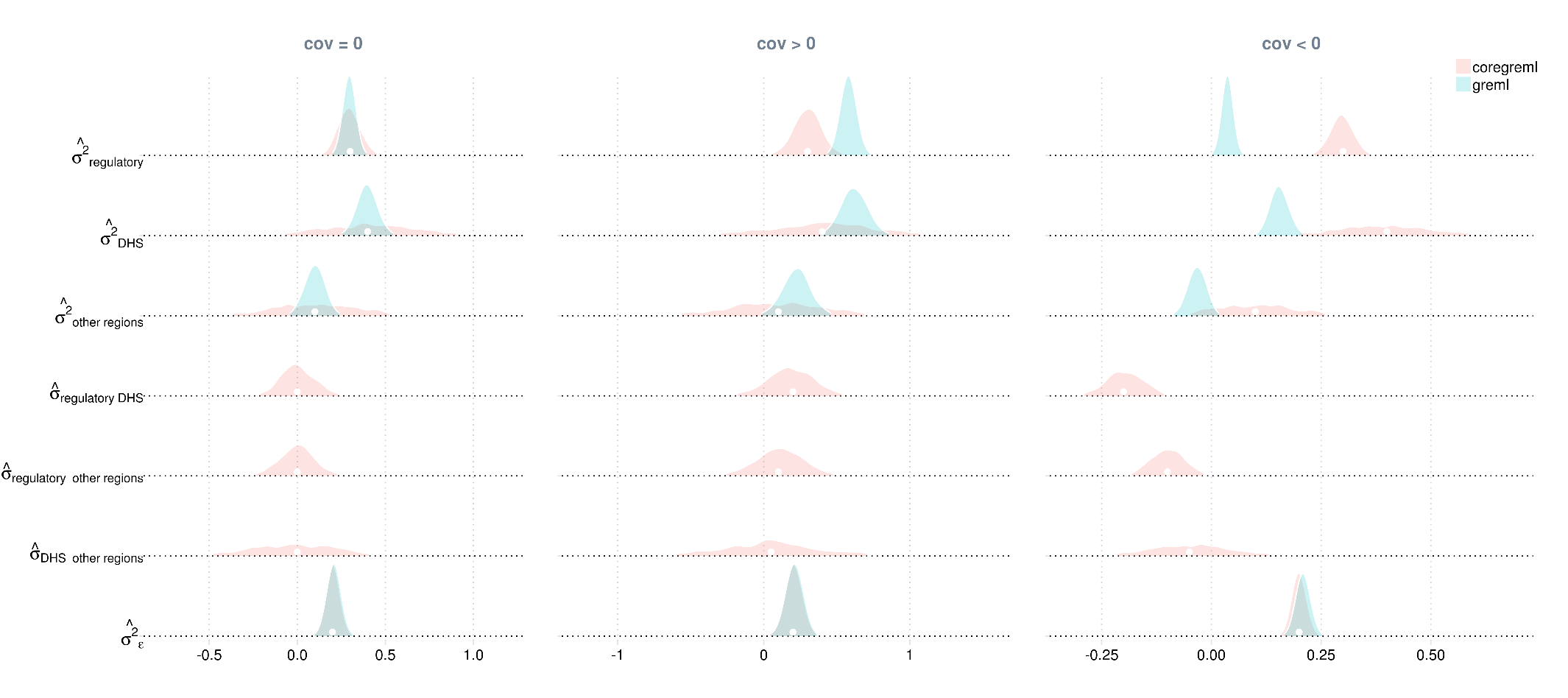

Supplementary Figure 3. Sampling distributions of model parameters by estimation method for genomic partitioning analysis. Five-hundred replicates of phenotypic data (n=10,000) were simulated under each of three parameter settings, where covariances between phenotypic effects of the three functional regions of the genome were zero (cov=0), positive (cov>0) and negative (cov<0). For each replicate, model parameters were estimated using the traditional method, i.e., GREML, and the proposed method, i.e., CORE GREML. True values of model parameters are in dots.

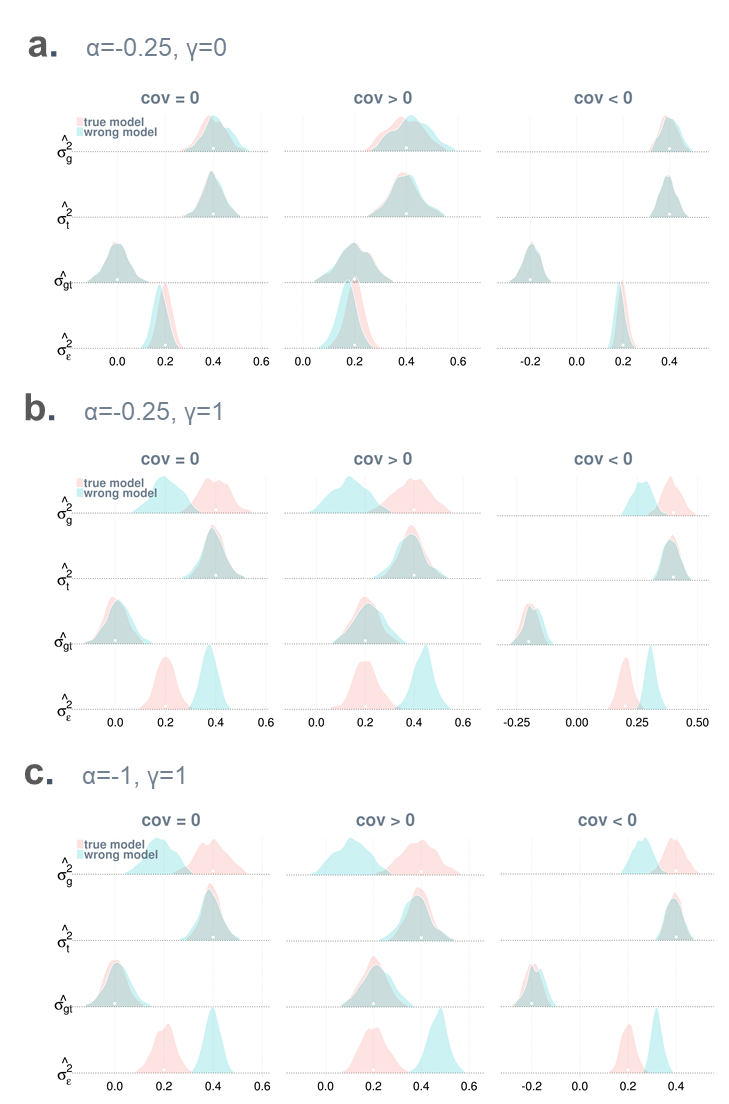

Supplementary Figure 4. Misspecification of genetic architecture in the estimation model affects variance estimates but not covariance estimates by CORE GREML. Shown in each panel are sampling distributions of model parameters by CORE GREML when the estimation model assumes the correct genetic architecture of the simulated trait (i.e., the true model is fitted) and when the estimation model mis-specifies the genetic architecture (i.e., a wrong model is fitted). The genetic architecture of the simulated trait is parameterised by linkage disequilibrium score, *w*, and minor allele frequency, *f*, in forms of $var(\beta_{i})\propto w_{i}^{\gamma}{[f_{i}(1-f_{i})]}^{1+\alpha}$ for any given causal SNP *i*, where *α* and *γ* control the extents to which w and *f* influence the variance of SNP-specific effects on phenotypes, i.e., var(β), respectively. Panels differ in values of *α* and *γ*, hence the genetic architecture of the simulated trait. Under each genetic architecture, there are three scenarios for the covariance between the phenotypic effects of the genome and those of the transcriptome, i.e., cov=0, cov>0 and cov<0. For each scenario, 500 replicates of phenotypes (each with n = 10,000) were simulated; the wrong estimation model always assumes *α* = -1 and *γ* = 0, while the true model assumes the values of *α* and *γ* identical to those of the simulation model. Regardless of estimation model, CORE GREML was applied for parameter estimation.

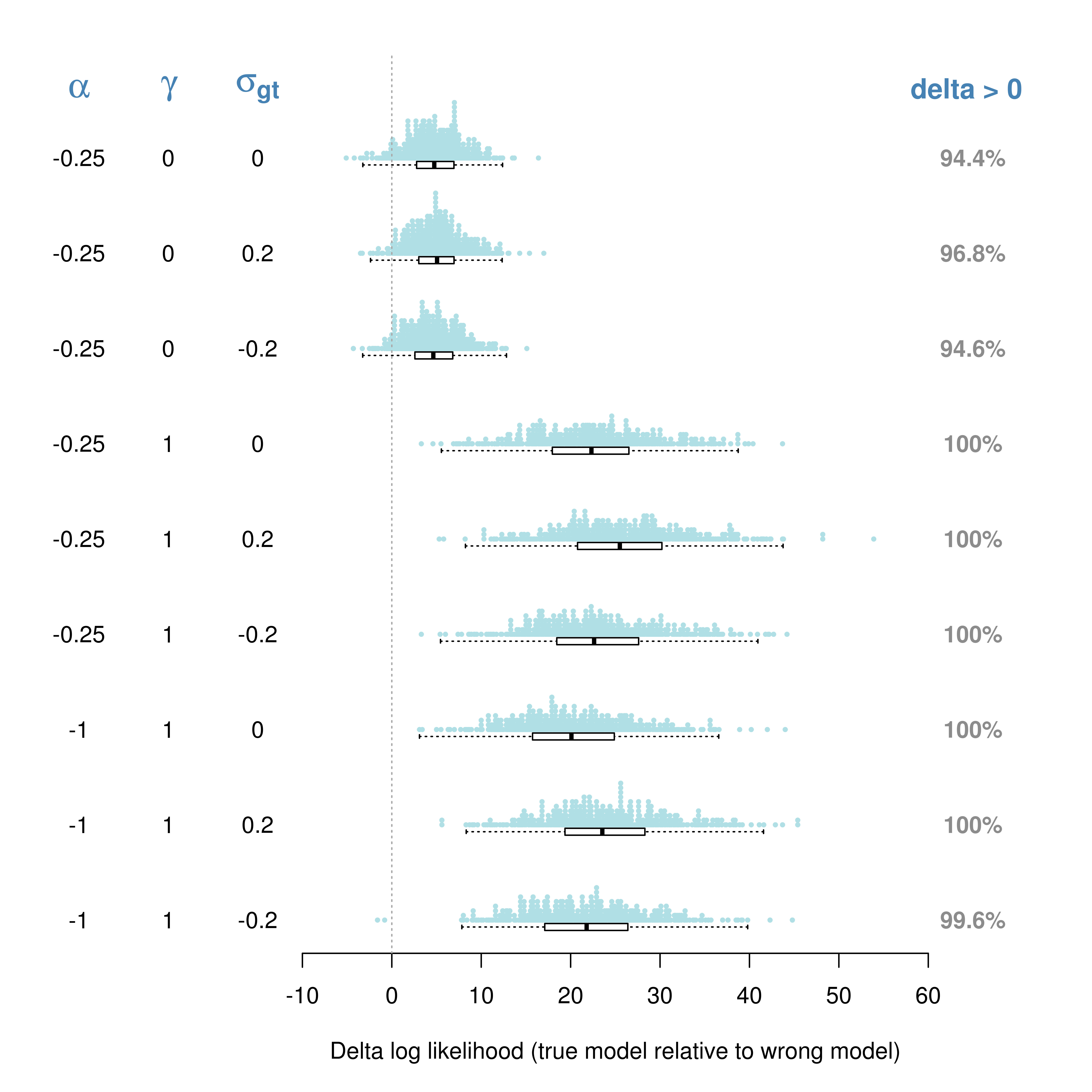
Supplementary Figure 5. Estimation model assuming the true genetic architecture of the simulated trait consistently shows a better fit than the model assuming a wrong genetic architecture. Shown in each row is the distribution of differences in log likelihood between the estimation model (true model) that assumes the correct genetic architecture of the simulated trait and a ‘wrong model’ that mis-specifies the genetic architecture. Irrespective of the estimation model, CORE GREML is used for parameter estimation. The difference in log likelihood is computed by subtracting the log likelihood of the wrong model from that of the true model, such that values above zero indicate that the true model has a better fit than the wrong model. The genetic architecture of the simulated trait is parameterised by linkage disequilibrium structure, *w*, and minor allele frequency, *f*, in forms of $var(\beta_{i})\propto w_{i}^{\gamma}{[f_{i}(1-f_{i})]}^{1+\alpha}$ for any given causal SNP *i*, where *α* and *γ* control the extents to which *w* and *f* influence the variance of the SNP-specific effects on phenotypes, i.e., var(β), respectively. Combinations of *α* and *γ* values give rise to different genetic architectures (information displayed in columns 1 & 2 on the left). Under each genetic architecture, there are three scenarios for the covariance between the phenotypic effects of the genome and those of the transcriptome (column 3 on the left). For each scenario, 500 replicates of phenotypes (each with n = 10,000) were simulated; the wrong estimation model always assumes *α* = -1 and *γ* = 0, while the true model assumes the values of *α* and *γ* identical to those of the simulation model.

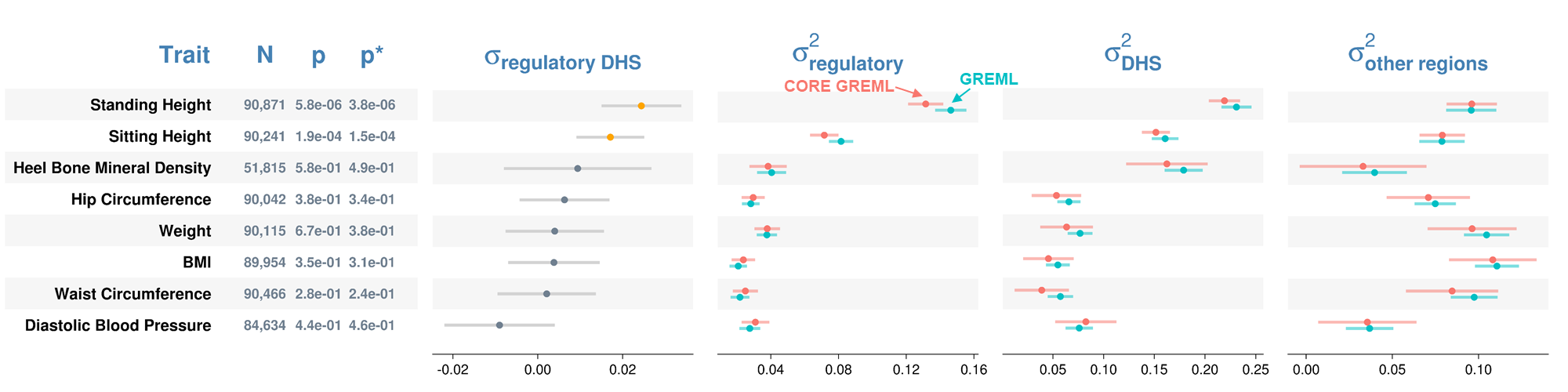

Supplementary Figure 6. Variance component estimates by method from genomic partitioning analyses. The three functional regions of the genome are regulatory regions, DHS, and all other regions. $\sigma_{regulatory DHS}$ denotes the covariance between genetic effects attributable to the regulatory regions and those attributable to the DHS on phenotypes. Error bars are 95% confidence intervals. p = p-values from likelihood ratio tests that compare GREML with CORE GREML to detect $\sigma_{regulatory DHS}$; p* = p-values from a sensitivity analysis where a rank-based inverse normal transformation was applied to phenotypic data to check the robustness of signals against the violation of the normality assumption held by both GREML and CORE GREML. Significant $\sigma_{regulatory DHS}$ are highlighted in orange. $\sigma_{\mathrm{regulatory}}^{2}$, $\sigma_{\mathrm{DHS}}^{2}$ , and $\sigma_{other regions}^{2}$ are phenotypic variances explained by the three functional regions. Residual variance estimates are omitted for simplicity. Fluid intelligence and years of education are excluded because either the phenotypic effects of the regulatory regions or those of the DHS were not significant.

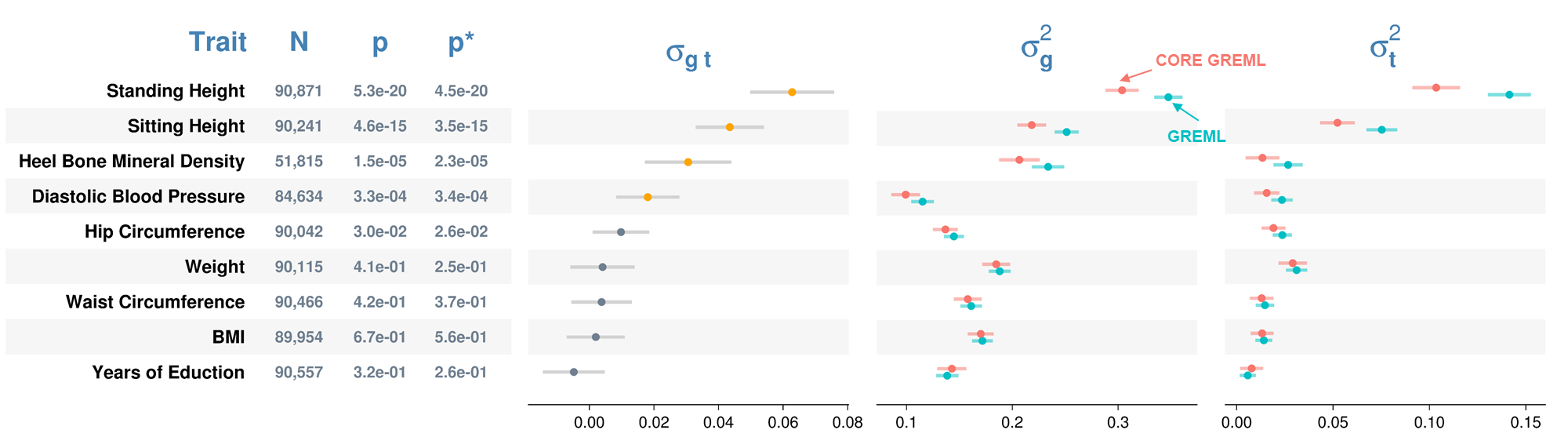

Supplementary Figure 7. Variance component estimates by method from genome-transcriptome partitioning of phenotypic variance. $\sigma_{g t}$ denotes the covariance between genetic effects and effects of imputed gene expressions on phenotypes. Error bars are 95% confidence intervals; p = p-values from likelihood ratio tests that compare GREML with CORE GREML to detect $\sigma_{g t}$; p* = p-values from a sensitivity analysis where a rank-based inverse normal transformation was applied to phenotypic data to check the robustness of signals against the violation of the normality assumption held by both GREML and CORE GREML. $\sigma_{g}^{2}$ and $\sigma_{t}^{2}$ denote the phenotypic variances explained by the genome and by the imputed transcriptome, respectively. Significant $\sigma_{g t}$ are highlighted in orange. Residual variance estimates are omitted for simplicity. Fluid intelligence is excluded because the phenotypic effects of the imputed transcriptome on this trait was not significant after Bonferroni correction.

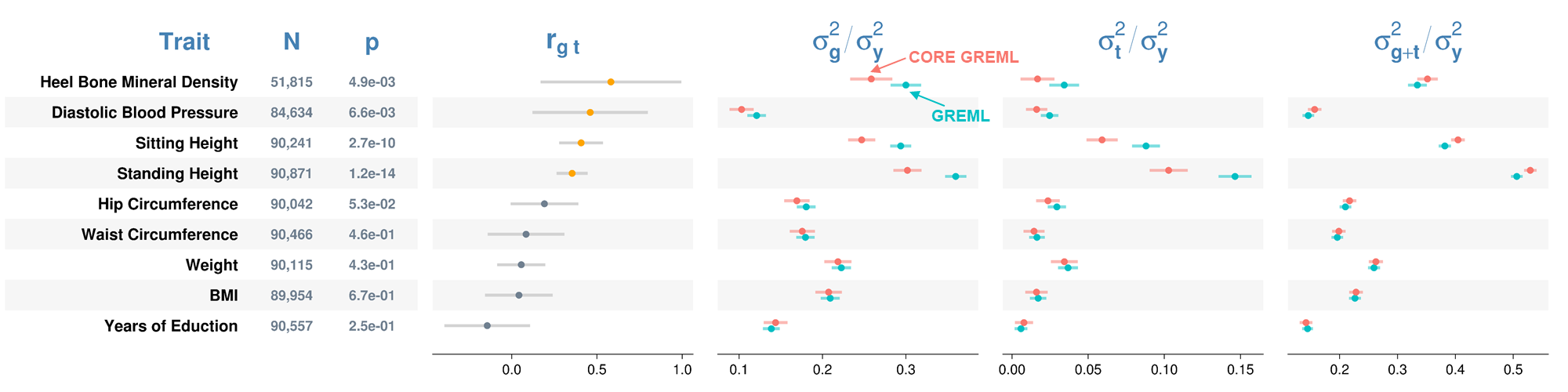

Supplementary Figure 8. Estimated proportions of phenotypic variance due to the genome and the imputed transcriptome by estimation method. $r_{g t}$ denotes the correlation between phenotypic effects of the genome and those of the imputed transcriptome. Error bars are 95% confidence intervals. P-values (p) are based on the Wald test statistic under the null hypothesis that $r_{g t}$ = 0, and significant correlations are highlighted in orange. Fluid intelligence is excluded because the phenotypic effects of the imputed transcriptome for this trait were not significant after Bonferroni correction.

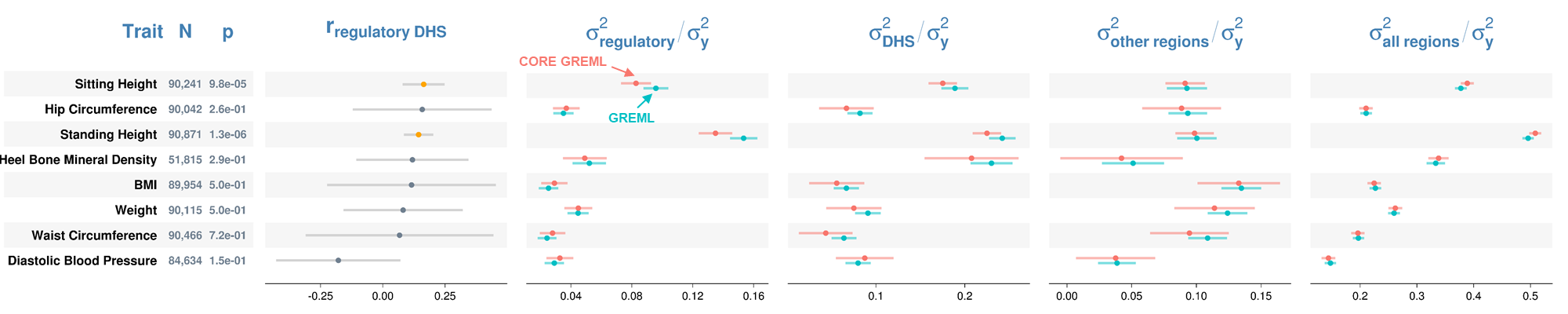

Supplementary Figure 9. Estimated proportions of phenotypic variance attributable to three functional genomic regions (i.e., heritability) by estimation method. The three functional regions are the regulatory regions, DHS, and all other regions. $r_{regulatory DHS}$ denotes the correlation between genetic effects of the regulatory region and DHS. Error bars are 95% confidence intervals. P-values (p) are based on the Wald test statistic under the null hypothesis that $r_{regulatory DHS}$= 0, and significant correlations are highlighted in orange. $\sigma_{\mathrm{regulatory}}^{2}$, $\sigma_{\mathrm{DHS}}^{2}$ , and $\sigma_{other regions}^{2}$ denote phenotypic variances explained by the three functional regions. $\sigma_{all regions}^{2}$= total genetic variance of all three functional regions; $\sigma_{y}^{2}$= total phenotypic variance. Fluid intelligence and years of education are excluded because either the phenotypic effects of the regulatory regions or those of the DHS were not significant.

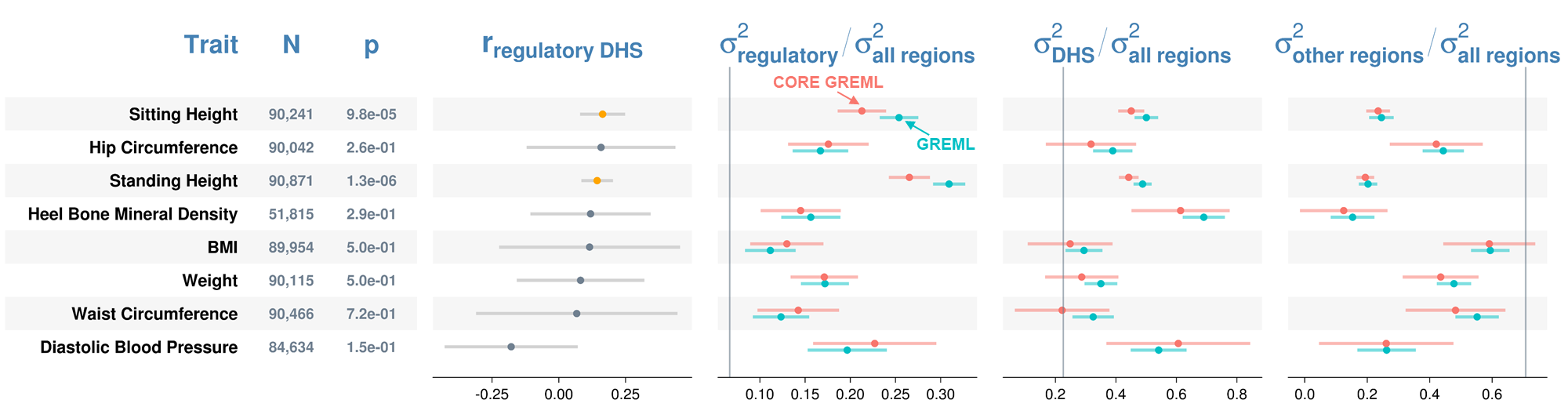

Supplementary Figure 10. Estimated proportions of total genetic variance attributable to three functional regions of the genome by estimation method. The three functional regions are the regulatory regions, DHS, and all other regions. $\sigma_{all regions}^{2}$= total genetic variance of all three functional regions; $r_{regulatory DHS}$ = correlation between genetic effects of the regulatory regions and DHS. Error bars are 95% confidence intervals. P-values (p) are based on the Wald test statistic under the null hypothesis that $r_{regulatory DHS}$ = 0, and significant correlations are highlighted in orange. Vertical lines are percentages of SNPs from the three functional regions, and conceptually, they are expected proportions of total genetic variance explained by the three functional regions of the genome assuming all genome-wide SNPs have an equal contribution to phenotypic variation. Fluid intelligence and years of education are excluded because either the phenotypic effects of the regulatory regions or those of the DHS were not significant.

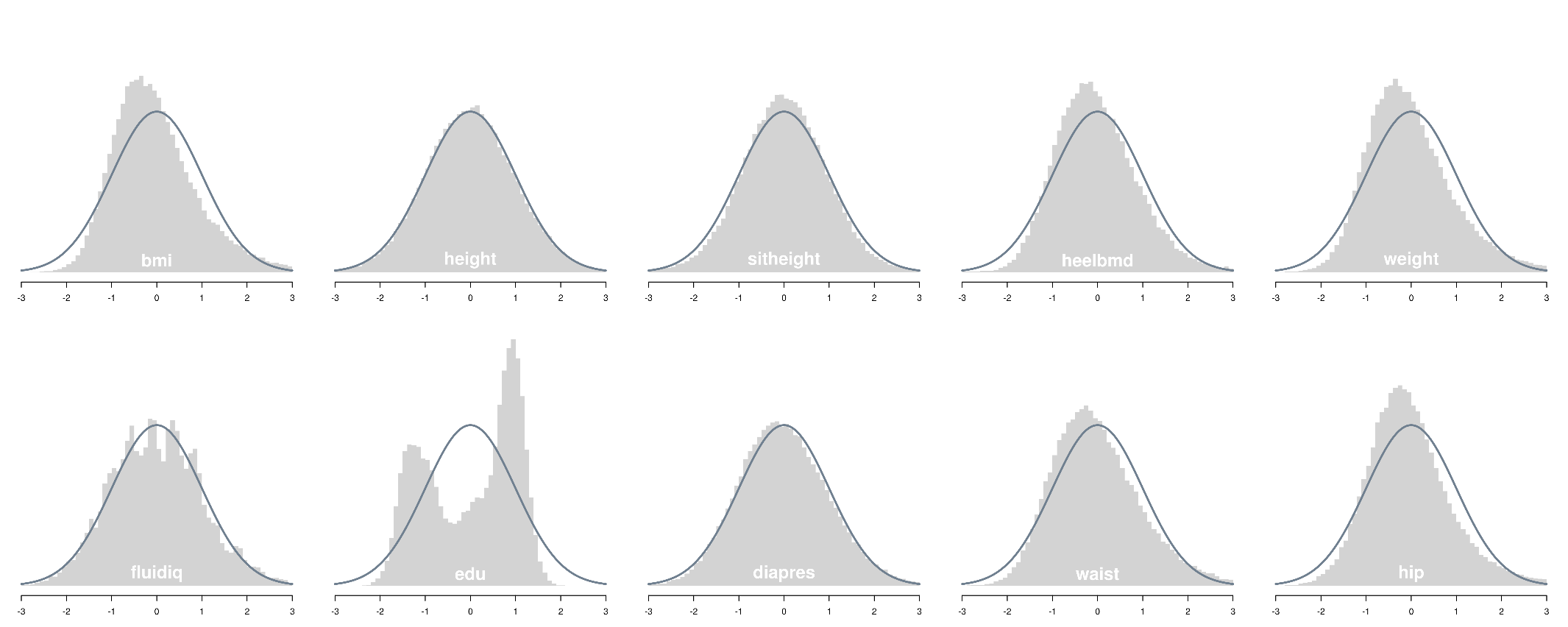

Supplementary Figure 11. Histograms of phenotypic data for ten selected traits from the UK Biobank. Traits from top left to bottom right are body mass index, standing height, sitting height, heel bone mineral density, weight, fluid intelligence, years of education, diastolic blood pressure, waist circumference and hip circumference. Data were prepared in three sequential steps: 1) adjustment for age, sex, birth year, social economic status, population structure, assessment centre, and genotype batch; 2) standardization; and 3) removal of data points outside +/- 3 standard deviations from the mean. The density function of Normal (0, 1) is superimposed as the reference to highlight deviations from normality.

Supplementary Table 1. True model parameter values for simulation models under three parameter settings.

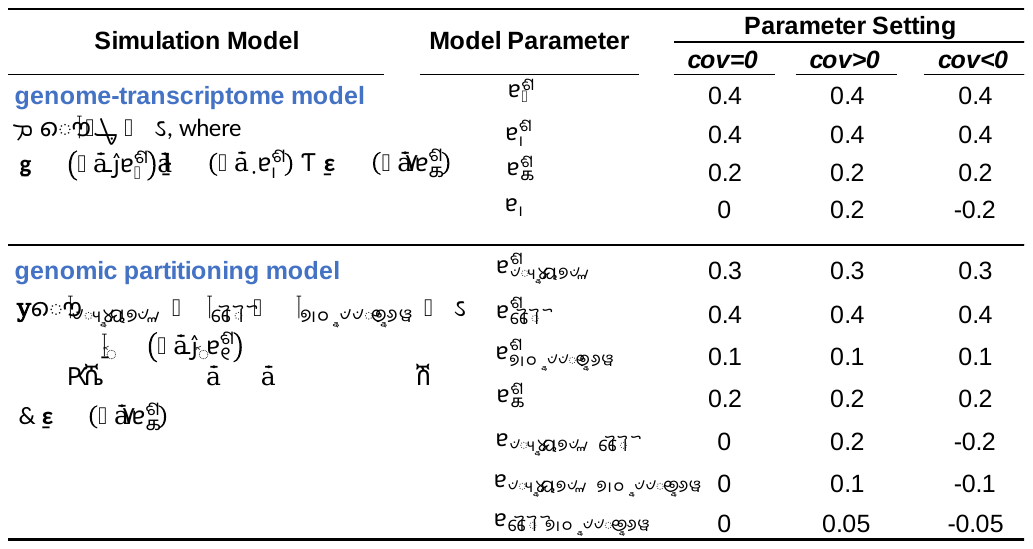

**A** & **T** are kernel matrices constructed using genotypes of 1,131,002 SNPs and using imputed expression levels of 227,664 genes collapsed across 43 tissues, respectively. **I** is an identity matrix. **A_i_** is the genomic relationship matrix constructed using SNPs from functional region *i* of the genome.

Supplementary Table 2. The number of genes for which expression levels were imputed across 43 non-sex-specific tissues.

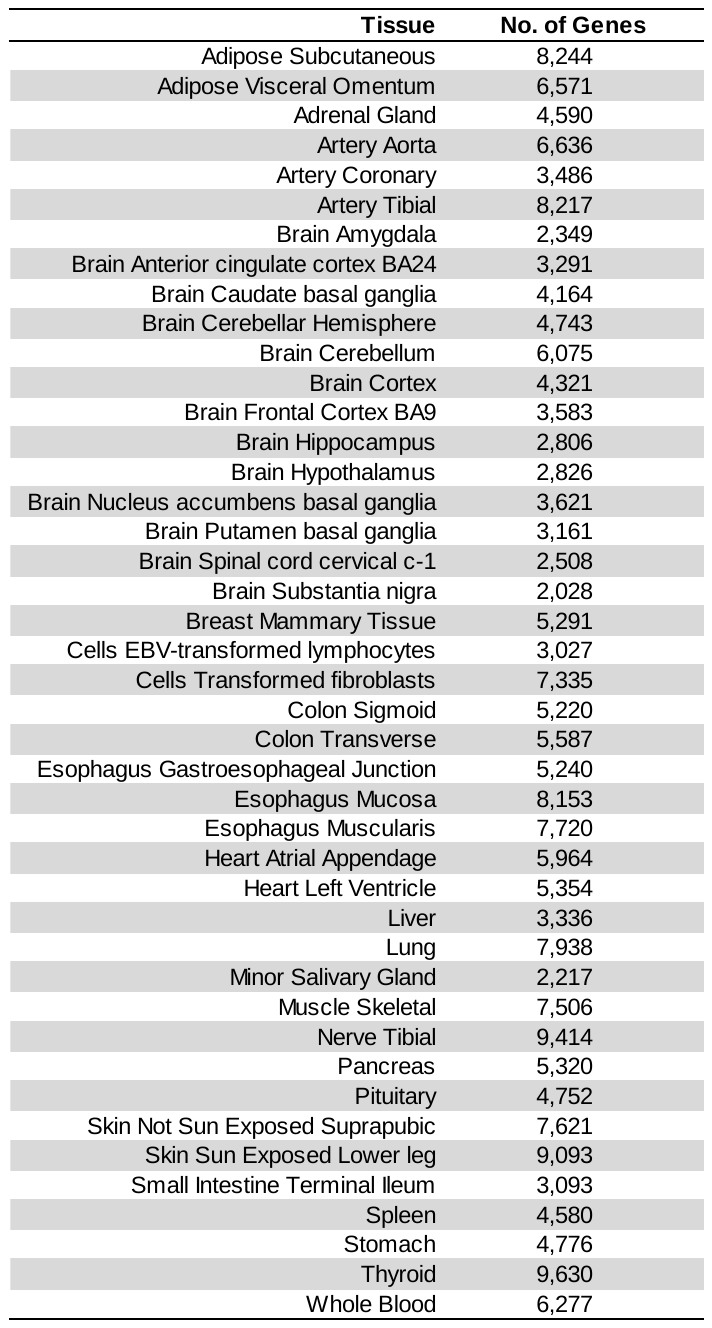

Note: Transcriptome imputation was based on GTEx V7 models (<http://predictdb.org/>).

Supplementary Table 3. Heritability estimates by estimation model.

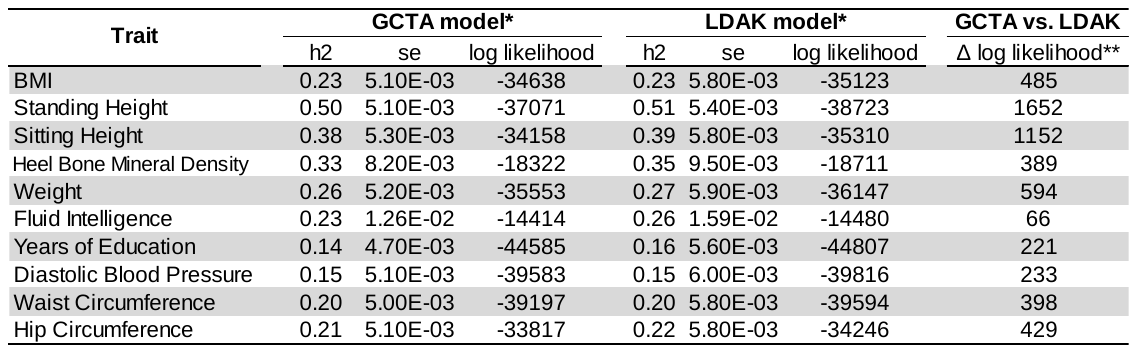

*Fitted GCTA model and LDAK model differ in the assumption about the variance of SNP-specific effects on phenotypes, $var(\beta)$. In general, it is assumed that for a given SNP *i*, the variance of its effects on phenotypes is proportional to its linkage disequilibrium score, *w*, and minor allele frequency, *f*, i.e.,$var(\beta_{i})\propto w_{i}^{\gamma}{[f_{i}(1-f_{i})]}^{1+\alpha}$, where parameter *γ* modulates the effect of *w* and *α* modulates the effect of *f* on $\mathrm{var}\left( \beta_{i} \right)$. In the GCTA model, *α* and *γ* were assumed to be -1 and 0, respectively; in the LDAK model, *γ* was assumed to 1 and *α* was set to the recommended default, -0.25 (Speed et al., 2017). Regardless of the estimation model, GREML is used for heritability estimation. **Δ log likelihood is derived by subtracting the log likelihood of the LDAK model from that of the GCTA model.

Supplementary Table 4. Likelihood of estimation models when GREML and CORE GREML were applied for the genome-transcriptome partitioning of phenotypic variance.

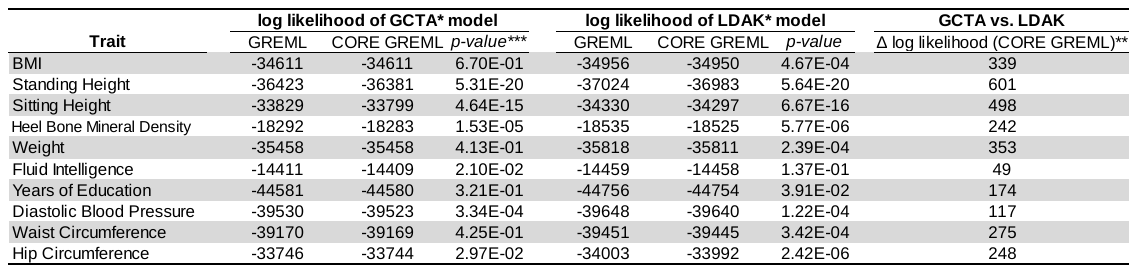

*Fitted GCTA model and LDAK model differ in the assumption about the variance of SNP-specific effects on phenotypes, $var(\beta)$. In general, it is assumed that for a given SNP *i*, the variance of its effects on phenotypes is proportional to its linkage disequilibrium score, *w*, and minor allele frequency, *f*, i.e.,$var(\beta_{i})\propto w_{i}^{\gamma}{[f_{i}(1-f_{i})]}^{1+\alpha}$, where parameter *γ* modulates the effect of *w* and *α* modulates the effect of *f* on$var(\beta_{i})$. In the GCTA model, *α* and *γ* were assumed to be -1 and 0, respectively; in the LDAK model, *γ* was assumed to 1 and *α* was set to the recommended default, -0.25 (Speed et al., 2017). **Δ log likelihood is derived by subtracting the log likelihood of the LDAK model from that of the GCTA model, when CORE GREML is used for parameter estimation for both models.***p-values are from likelihood ratio tests that compare GREML with CORE GREML to detect the covariance between random effects of the genome and those of the transcriptome on phenotypes.

Supplementary Table 5. GREML estimates of genetic variance partitioned by functional genomic region.

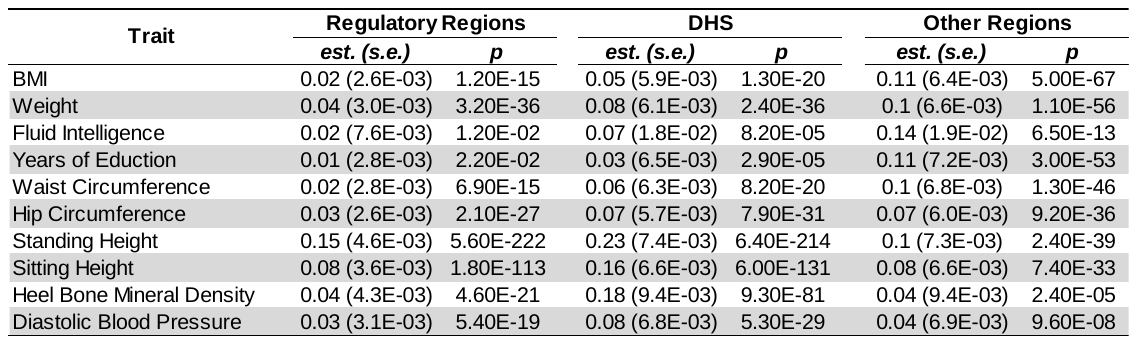

Note: p-values are based on the Wald test statistic with one degree of freedom under the null hypothesis that the variance component of interest is zero.

Supplementary Table 6. Correlations between off-diagonal entries of kernel matrices used for variance-components estimation.

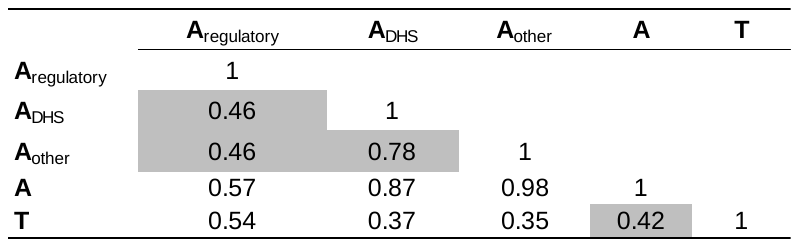

**A**, **A**_regulatory_, **A**_DHS_ and **A**_other_ are genomic relationship matrices constructed using available SNPs from the entire genome, the functional regions, the DHS and all other regions, respectively. **T** is a kernel matrix constructed using the imputed expression levels of genes collapses across 43 tissues. Note all matrices are based on data from 10,000 UK biobank participants randomly selected for our simulations.

**Supplementary Note**

In this note, we report how well GREML recovers phenotypic effects of the imputed transcriptome. We simulated 500 replicates of phenotypic data using the genome-transcriptome model (Table 1 of main text) under settings with and without a random term for the phenotypic effects of the imputed transcriptome ($\sigma_{t}^{2}$= 0 vs. $\sigma_{t}^{2}$= 0.4). For both settings, the covariance between phenotypic effects of the genome and those of the imputed transcriptome was set to zero. For each replicate, we fitted two models, a ‘G model’ that breaks phenotypic effects into effects of the genome and residuals, i.e., y=g+ε, and a ‘G-T model’ that decomposes phenotypic effects into effects of genome and of the imputed transcriptome and residuals, i.e., y=g+t+ε; both models were estimated using GREML. We declared the presence of a transcriptomic effect when the G-T model had a better fit than the G model via a likelihood ratio test with one degree of freedom. We found that the type I error rate was controlled (0.042) under the zero transcriptomic effect setting. Under both settings, G-T model yielded unbiased estimates of model parameters (Sup.N. Fig. 1 a). On the other hand, the G model produced unbiased estimates only under the null setting, as expected. Interestingly, in the presence of a transcriptomic effect, the estimated genetic variance by the G model was equivalent to the sum of estimated variances due to the genome and imputed transcriptome by the G-T model (Sup.N. Fig. 1 a), such that the phenotypic variance explained by the two models were similar. This indicates that SNP-based heritability estimates by GREML can be inflated by phenotypic variance due to the transcriptome. Importantly, although both models captured a similar amount of phenotypic variance, the fit of the G-T model was far better than that of the G model (Sup.N. Fig. 1 b), indicating the partition of phenotypic variance represented by the G-T model is closer to the truth than that represented by the G model.

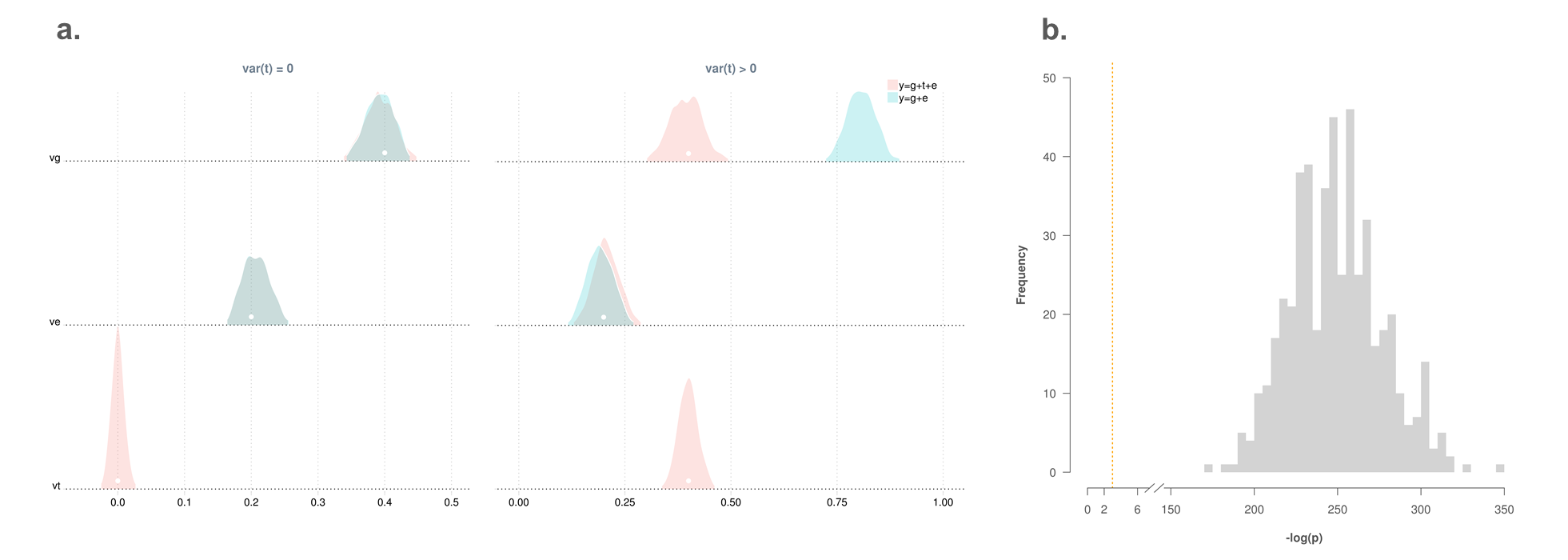

Sup.N.Figure 1. GREML recovers phenotypic effects of the imputed transcriptome. Five-hundred replicates of phenotypic data (n=10,000) were simulated under settings without and with a term for phenotypic effects of the imputed transcriptome, denoted as var(t)=0 and var(t)>0, respectively. For each replicate, two linear mixed-effects models were fitted, one that did not include a term for the phenotypic effect of the imputed transcriptome, i.e., y=g+ε, and the other did, i.e., y=g+t+ε. All model parameters were estimated using GREML. **Panel a**. estimated density of model parameters. True values are in dots. **Panel b**. histogram of p-values from likelihood ratio tests (df=1) that compared the two models for replicates simulated under var(t)>0. P-values are log transformed with the statistical significance threshold, i.e., -log (0.05), being indicated by the vertical line. Values above the threshold indicate that y=g+t+ε had a better fit than y=g+ε.
